# Quantifying the capacity for assisted migration to achieve conservation and forestry goals under climate change

**DOI:** 10.1101/2022.03.29.486208

**Authors:** Yibiao Zou, Gregory A. Backus, Hugh D. Safford, Sarah Sawyer, Marissa L. Baskett

**Author notes:** Correspondence: Yibiao Zou.

## Abstract

Many tree species might be threatened with extinction because they cannot disperse or adapt quickly enough to keep pace with climate change. One potential, and potentially risky, strategy to mitigate this threat is assisted migration, the intentional movement of species to facilitate population range shifts to more climatically suitable locations under climate change. The ability for assisted migration to minimize risk and maximize conservation and forestry outcomes depends on a multi-faceted decision process for determining, what, where, and how much to move. To quantify how the benefits and risks of assisted migration could affect the decision-making process, we used a dynamical vegetation model parameterized with 23 tree species in the western United States. We found that most of the modeled species are likely to experience a substantial decline in biomass, potentially facing regional extinction by 2100 under the high-emission SSP5-85 climate-change scenario. Though simulations show assisted migration had little effect on the forestry goal of total biomass across all species, its effects on the conservation goal of promoting individual species’ persistence were far more substantial. Among eight assisted migration strategies we tested that differ in terms of life cycle stage of movement and target destination selection criteria, the approach that conserved the highest biomass for individual species involved relocating target seedlings to areas with the highest canopy openness. Although this strategy significantly reduced extinction risk for six at-risk species compared to no action, it also slightly reduced biomass of four species, due to increasing competition. Species with relatively weak tolerance to drought, fire or high temperature were the most likely candidate groups for assisted migration. This model framework could be applied to other forest ecosystems to evaluate the efficacy of assisted migration globally.

## Introduction

Climate change imposes extinction risk on species that cannot keep pace through acclimatization, adaptation, or dispersal (Davis Margaret & Shaw Ruth, 2001; Holt, 1990). Of particular concern are foundational species with limited dispersal ability such as forest trees, which can lead to substantial ecosystem-wide consequences if they cannot keep pace with climate change (Kreyling et al., 2011). Climate change has already placed stress on forests globally by increasing the extremity of droughts, changing fire regimes, and amplifying climate variability (Williams & Dumroese, 2013). Results of global climate models and species distribution models suggest that climatically suitable habitat for many tree species will contract or shift to higher latitudes or altitudes in the coming decades (Coops & Waring, 2011; Iverson & McKenzie, 2013). Given the rapidity of current climate change, most tree species face the risks of range decline or extinction because they are unlikely to track climate shifts through natural dispersal alone or to develop novel adaptations (Nathan et al., 2011; Williams & Dumroese, 2013; Zhu et al., 2012).

One potential strategy to preserve endangered tree species, forest biodiversity and ecosystem services under climate change is assisted migration (AM; also known as managed relocation), the intentional movement of species to more suitable locations outside their current range given projected climate change (McLachlan et al., 2007; Williams & Dumroese, 2013). Although AM might increase the likelihood of persistence for vulnerable species under climate change, its potential risks have generated scientific controversy and debate (Hewitt et al., 2011; Minteer & Collins, 2010; Ricciardi & Simberloff, 2009; Richardson et al., 2009). One risk of AM is establishment failure from relocating individuals to the wrong place, at the wrong time, or from the wrong source location, potentially stressing the source population without establishing a new self-sustaining population (Kreyling et al., 2011). Another risk of AM is negative effects relocated populations can have on the recipient community, such as is the case with invasive species. Typical invasive-type impacts include spreading diseases or parasites, negative interspecific interactions (such as competition or predation), and hybridization (Ricciardi & Simberloff, 2009; Simler et al., 2019).

Assisted migration in forest systems might be employed to acheive either forestry or conservation goals, or potentially both (Pedlar et al., 2012). Forestry goals typically focus on forest productivity and ecosystem services, and conservation goals typically focus on species persistence, with diversity potentially connecting to both goals (Williams & Dumroese, 2013). Because AM is a single-species rather than a community-level management approach (Lawler & Olden, 2011), AM is likely to increase persistence of vulnerable species and promote conservation goals. However, if only a small number of species are at risk of extinction from climate change, a single-species approach like AM could theoretically be effective as a community-level management approach (Backus et al., *in press*). Whether or not a single-species approach like AM supports forestry goals of productivity, ecosystem services and diversity depends on the community-level outcomes. Understanding this capacity depends on evaluating how relocated populations compete with non-target species and contribute to overall system productivity.

Achieving both conservation and forestry goals through AM, as well as reducing its risks, depends on a multi-faceted decision process relating to which species, when, where, and how much to move. For the question of which species to move, in the face of a warmer, drier future climate with more fire, we expect that tree species with poor dispersal capacity and weak tolerance to warm temperature, drought or fire would be at risk of population decline or extinction, and therefore potential candidate target species for AM (Clark et al., 2016; Williams & Dumroese, 2013). For the question of when to move within a species’ life cycle, the current common practice is moving seedlings cultivated in the greenhouse (Haase et al., 2019). An alternative approach of directly moving seeds to recipient communities can reduce the logistical, financial, and time costs of seedling cultivation but might reduce relocated species germination and survival through earlier life stages.

For the question of where to move, there are several criteria for choosing target locations. Targeting the closest climatically suitable locations to the species’ historical range could reduce the risk of introducing novel competition to species already extant in the recipient community (Pedlar et al., 2012). Another strategy is to select the locations with low canopy cover as to reduce light and water competition and increase establishment success for AM individuals. On one hand, moving target species to sites which recently experienced fire can take advantage of fire’s reduction of both canopy cover and understory cover, as well as the postfire abundance of bare mineral soil (Welch et al., 2016). On the other hand, moving species to sites with the least expected fire frequency can reduce the potential for fire disturbance to affect establishment success. For the question of how much to move, increasing the number and frequency of AM actions might increase the likelihood of establishment of the target species, and therefore AM benefits, and enhance the risk of competition for non-target species. Quantifying these different aspects for implementing AM requires building on existing SDM approaches to create a dynamic modeling framework (Iverson & McKenzie, 2013) that incorporates dispersal and competition dynamics. Incorporating dispersal dynamics can determine whether AM can out-perform expectations under natural levels of dispersal, while accounting for competition is necessary for understanding its role in the relative performance of the different approaches to selecting target locations and different AM intensities.

Here we use a dynamic vegetation model (DVM) to quantify how forestry and conservation outcomes of AM depend on different decision-making approaches for which species, when, where, and how much to move. We parameterized the model with 23 dominant tree species from forest ecosystems in the North American Mediterranean Climate Zone (NAMCZ, Safford et al., 2021) and identified target species traits based on projected population declines below a threshold value under future climate change. We considered four types of destination site-selection for AM: **m**inimum-**d**istance destinations (MD), **l**east-**c**ompetition destinations (LC), **p**ost-**f**ire destinations (PF) and **l**east-**f**ire destinations (LF). Additionally, we incorporated two AM types differed by relocated life stage: see**d A**M (DA) and seedlin**g A**M (GA). We also explored a range of values for AM intensity in terms of its frequency, its duration, the number of locations targeted, and the number of individuals moved. Simulating climate change and different AM management strategies, we measured forestry goals in terms of total biomass, conservation goals in terms of individual species biomass, persistence, and both with biomass-weighted gamma diversity.

## Materials and methods

### I. Study region

We simulated forest dynamics in the mountain landscapes of the coastal Mediterranean-climate western United States, much of which is charactierized by a Mediterranean-type climate (Safford et al., 2021). In this region, climate change is increasing the frequency, magnitude, and duration of drought events, which also enhances the intensity and frequency of wildfires (Mann & Gleick, 2015; Miller et al., 2009). Several tree species in this region face the threat of range contraction or local extinction under projected increases in frequency and severity of fire and drought events, making them strong candidates for AM (Loarie et al., 2008; Rogers et al., 2011). Among these AM candidate species are economically important species such as ponderosa pine (*Pinus ponderosa*) and currently rare species like foxtail pine (*Pinus balfouriana*) (Richardson et al., 2009). Because many of these species take decades to reach maturity (Bonner et al., 2008), urgent conservation might be necessary to prevent extinction and losses of ecosystem services (Stephens et al., 2020).

We focused on a region made up of two connected line segments, 1 degree longitude in width, following the Sierra Nevada (36°11’ N, 119°3’ W to 41°23’ N, 122°47’ W) and the Cascade Range (41°23’ N, 122°47’ W to 48°36’ N, 121°39’ W). We divided this area into 1560 5-arcmin^2^ grid cells (12 wide and 130 long), each consisting of 200 patches (833 m^2^ each) in which we simulated forest dynamics representing the mean dynamics of the entire cell (Gutiérrez et al., 2016). While the spatial location of grid cells is explicit, the spatial structure of patches is implicit. In the model simulation, we simplified the spatial structure of the study region into a 130×12 matrix, where the distance between the center of any two Von Neumann neighbor grid cells is 5-arcmin (i.e., ∼10 km).

At this 5-arcmin resolution scale, we used digital Little’s maps to estimate initial tree species occurrence (Little, 1971), the LEMMA dataset for determining initial tree basal area (Ohmann & Gregory, 2002), and the WorldClim version-2.1 dataset for elevation data and climate data (Fick & Hijmans, 2017). We modeled 23 dominant tree species from the Sierras and Cascades with accessible physiological parameters (**Table 1**).

**Table 1:** Species-specific parameter set of 23 tree species used in this study. . Acronyms of columns of traits: minimum growing degree-day requirement (*kDDMin_s_*, °C), drought tolerance (*kDrtol_s_*), fire tolerance class (*kFi_s_*; as class number increases, fire tolerance decreases), minimum and maximum winter temperature tolerances (*kWiTN_s_* and *kWiTX_s_*, °C). See S2 Parameterization in the appendix for parameter value sources, justification, and calculations.

| *Scientific name* | *Common name* | *Species_id* | *kDDMin<sub>s</sub>* | *kDrTol<sub>s</sub>* | *kFi<sub>s</sub>* | *kWiTN<sub>s</sub>* | *kWiTX<sub>s</sub>* |
| --- | --- | --- | --- | --- | --- | --- | --- |
| *Abies amabilis* | Pacific silver fir | 1 | 562 | 0.17 | 1 | -5.43 | 3.66 |
| *Abies grandis* | Grand fir | 2 | 594 | 0.28 | 2 | -5.24 | 4.28 |
| *Abies lasiocarpa* | subalpine fir | 3 | 526 | 0.23 | 1 | -5.71 | 1.39 |
| *Abies procea* | Noble fir | 4 | 601 | 0.23 | 2 | -4.01 | 3.07 |
| *Acer macrophyllum* | Big-leaf maple | 5 | 1010 | 0.54 | 2 | -1.24 | 8.3 |
| *Arbutus menziesii* | Madrone | 6 | 868 | 0.58 | 2 | -2.56 | 8.54 |
| *Chamaecyparis*<br>*nootkatensis* | Alaska cedar | 7 | 540 | 0.21 | 1 | -5.65 | 2.26 |
| *Picea engelmannii* | Engelmann spruce | 8 | 536 | 0.28 | 1 | -5.66 | 2.4 |
| *Pinus contorta latifolia* | Lodgepole pine | 9 | 500 | 0.37 | 2 | -5.56 | 3.94 |
| *Pinus monticola* | Western white pine | 10 | 595 | 0.38 | 2 | -5.07 | 3.92 |
| *Pinus ponderosa* | Ponderosa pine | 11 | 817 | 0.57 | 4 | -4.67 | 7.09 |
| *Pseudotsuga menziesii*<br>*menziesii* | Pacific coast Douglas-fir | 12 | 659 | 0.48 | 3 | -5.01 | 5.94 |
| *Quercus garryana* | Oregon white oak | 13 | 1132 | 0.52 | 4 | -1.01 | 6.71 |
| *Tsuga heterophylla* | Western hemlock | 14 | 623 | 0.19 | 1 | -4.98 | 4.32 |
| *Tsuga mertensiana* | Mountain hemlock | 15 | 504 | 0.32 | 1 | -5.66 | 1.68 |
| *Abies concolor* | White fir | 16 | 697 | 0.49 | 2 | -3.54 | 5.66 |
| *Abies magnifica* | Red fir | 17 | 604 | 0.44 | 3 | -4.11 | 3.54 |
| *Calocedrus decurrens* | Incense cedar | 18 | 880 | 0.52 | 3 | -2.67 | 6.42 |
| *Pinus jeffreyi* | Jeffrey pine | 19 | 444 | 0.49 | 3 | -5.63 | 4.13 |
| *Pinus lambertiana* | Sugar pine | 20 | 859 | 0.54 | 3 | -2.77 | 6.66 |
| *Pinus albicaulis* | Whitebark pine | 21 | 361 | 0.36 | 2 | -6.06 | 0.69 |
| *Quercus kelloggii* | California Black oak | 22 | 1035 | 0.57 | 3 | -1.59 | 8.21 |
| *Pinus balfouriana* | Foxtail Pine | 23 | 270 | 0.42 | 4 | -7.7 | 2.86 |

### II. Components of the model

Our model simulates forest dynamics of individual trees, climate change, and management actions in R-3.6.2. The model has four sub-models that change over each one-year time step: demographic vegetation, climate, fire, and assisted migration (Figure 1). The following sections explain these sub-models.

**Figure 1:**
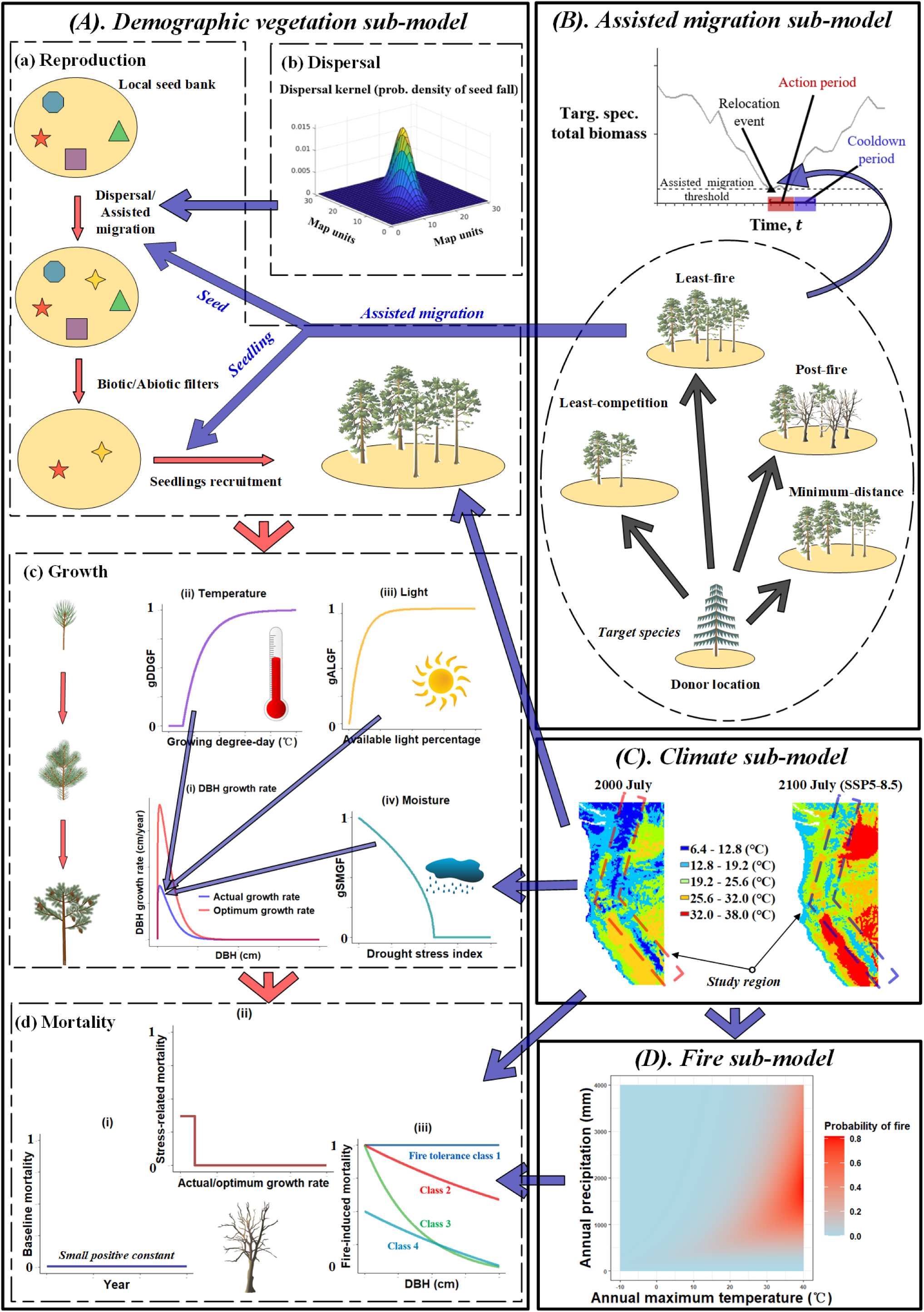
Model diagram. The model has four components: demographic vegetation, assisted migration, climate, and fire. Blue influencing arrows indicate connections between these components. (A) The demographic vegetation sub-model simulates life cycle that each individual tree steps through in each time step as indicated by the red arrows: (A.a-b) reproduction & dispersal, (A.c) growth, and (A.d) mortality. For (A.a) reproduction, there is a local seed bank, dispersal can stochastically add new species to the seed bank, local climate or competition can prevent some species at the seed bank from establishing as seedling, and the remaining species will end up growing. For (A.b) dispersal, seeds disperse follow a long-distance wind-driven dispersal kernel. The seed source is at the center of the X-Y plane, and the Z axis represents the probability density for seeds from source to fall in position (x, y). For (A.c) growth, seedlings or adult trees grow in the form of increased diameter at breast height (*DBH*) and height. Their growth depends on limiting index of (A.c.ii) growing degree-days, namely temperature (gDDGF), (A.c.iii) light availability (gALGF) and (A.c.iv) soil moisture (gSMGF), which determine the realized growth rate relative to the optimum growth rate (A.c.i). For (A.d) mortality, all stages of trees experience three sources of mortality: (A.d.i) baseline: each year a tree will experience a constant mortality probability; (A.d.ii) stress-related if a tree’s actual growth rate is below 0.1 of its optimum growth rate due to stress; (A.d.iii) fire-induced when a fire occurs, dependent on a tree’s fire tolerance class and *DBH*. The target species for AM also experiences (B) assisted migration (seed or seedling AM) when their biomass falls behind the AM threshold, which adds to (A.a) reproduction. For a target species, one AM action will continue a certain amount of years (action period), followed by a cooldown period during which no AM on the same species is implemented. We consider four characteristics that might determine the AM destination location: least fire (LF), least competition (LC), minimum distance (MD), and post-fire (PF). All of the four life history processes (A.a-d) in the (A) demographic vegetation sub-model directly or indirectly depends on basic climatic variables (temperature, precipitation and wind speed, etc.) and their derivative variables (growing degree-day, drought index, etc.), calculated through the (C) climate sub-model. The climate data also influences the fire probability in each grid cell, in the (D) fire sub-model, which affects (A.d) mortality.

#### (A). Demographic vegetation sub-model

For the demographic sub-model, we modified the PLANT sub-model of ForClim v.3.0 (Gutierrez et al., 2016) by adding dispersal (Figure 1A). Many previous studies validated several versions of ForClim by predicting reasonable forest species composition and productivity across a wide range of climate gradients across the world (Bugmann & Solomon, 2000), including the western United States (Bugmann & Solomon, 2000; Gutiérrez et al., 2016).

The PLANT sub-model of ForClim simulates forest demographics within each independent patch, where individual tree demographics (recruitment, growth, and mortality) follow three independent functions of biotic interaction and abiotic constraints. First, in the recruitment function, the model adds new seedlings to a patch based on the seeds available in the local species pool and filtered by species-specific responses to minimum winter temperature, growing degree-days, and light availability on the forest floor (see S1.1.1 Tree reproduction). We assumed species with basal area larger than a threshold *T_p_* in a patch have effective seed bank in that patch to support their presence. Here basal area is a surrogate for biomass (linear approximation, see Eqs (29) & (30) in Appendix), which is more commonly used to determine species’ presence status (Chang & Bourque, 2020). In the growth function, each individual living tree in the model has a chance to increase in diameter and biomass (see S1.1.2 Tree growth). The optimum tree growth rate mainly depends on net carbon assimilation, following the carbon budget approach by Moore (1989), while light availability across the crown, growing degree-days (GDD) and drought stress limit the realized growth rate. In the mortality function, each individual living tree can experience mortality based on three mechanisms: baseline, stress-induced, and fire-induced (see S1.1.3 Tree mortality). Individuals experience mortality as a binomial draw weighted by the mortality probability. In this Demographic Vegetation sub-model, we only modify the recruitment function by introducing a dynamic local seed bank based on a two-dimensional long-distance seed dispersal, while the other two functions remain unchanged from the base ForClim model (Bugmann & Solomon, 2000; Rasche et al., 2012).

#### (B). Assisted migration sub-model

In the Assisted Migration sub-model (Figure 1B), we simulate the relocation of seeds or seedlings of a species to a new location if that species’ biomass falls below a threshold value relative to its initial condition *T_AM_* (see Table 2). We set up the general framework of this sub-model based on the model developed in Backus & Baskett (2021). During any simulation in which a population of any given species falls below the threshold (see S1.2), we simulate an AM action to preserve this species. These AM actions repeat for *I_a_* years (years per relocation), followed by a cooldown period with a minimum of *I_c_* years between relocation event such that there is no AM for this species (Backus & Baskett, 2021). The cooldown period allows the AM species to establish to determine if AM promoted species persistence before taking further AM action.

**Table 2:** non-species-specific parameters used in the model, symbols used to represent them, values used, and units of these parameters. **See S1.1 Demographic vegetation sub-model** in the Appendix for more detailed description and sources of these parameters.

| Parameter | Symbol | Values | Units |
| --- | --- | --- | --- |
| Maximum number of newly established seedlings of one species in one patch | $N_{max}$ | 5 | individuals |
| Minimum basal area threshold for tree species presence | $T_p$ | 0.1 | m<sup>2</sup>/ha |
| Mean wind speed across our study region | $u$ | 5.689 | m/s |
| Standard deviation of the vertical velocity of the air | $\sigma$ | 0.25 | (m/s)<sup>2</sup> |
| Turbulence coefficient used to calculate dispersal kernel | $\kappa$ | 0.4 | - |
| AM threshold (percentage of initial species-specific biomass) | $T_{AM}$ | 30 | % |
| Number of simulated patches used to represent one grid cell | $N_{pa}$ | 200 | - |
| Area of each patch | $A_{pa}$ | 833 | m<sup>2</sup> |
| Years per relocation | $I_a$ | 2, 4, ..., 10 | years |
| Minimum years between relocation | $I_c$ | 4, 8, ..., 20 | years |
| Number of target grid cells for AM | $N_{gc}$ | 3, 6, ..., 15 | grid cells |
| Number of seedlings of the target species moved to one patch during seedling AM | $N_{sd}$ | 80, 160, ..., 400 | individuals |

To estimate suitable target sites for the target species under future climate conditions, we selected grid cells in which the projected bioclimatic conditions in 2100 were within the species’ climatic tolerance range. During each year within the relocation period, we set the number of target grid cells for AM as *N_gc_*. We simulated AM of two life stages in this sub-model, namely, seed and seedling. We simulated seed assisted migration (DA) by modifying the seed bank (relative recruitment) of the AM target species in the target grid cell, which gives the target species a relative recruitment advantage, namely a 30 times higher recruitment rate than other present species (even if the target species was not already present in the seed bank). For seedling assisted migration (GA), we based our simulation on the common practice in forestry of cultivating seedlings in a greenhouse for several years (Haase et al., 2019) before moving them to a target location. In our model, we simulated this by directly moving a certain number (*N_sd_* x 200) of seedlings of the target species with *DBH* = 1.27 *cm* (average size of cultivated seedlings among different tree species (Sáenz-Romero et al., 2021)) into a target grid cell, omitting the greenhouse cultivation processes. The number of seedlings moved to one patch was *N_sd_*, while 200 is the number of patches within one grid cell. During AM period, this sub-model simulates seed or seedling AM on all *N_gc_* target grid cells.

Within the potential target range determined by the anticipated future suitable climate, we considered four types of destination site-selection for AM: minimum-distance destinations (MD), least-competition destinations (LC), post-fire destinations (PF) and least-fire destinations (LF). Minimum-distance destinations are the target grid cells within the species’ potential 2100 range that are closest to the current range of the target species. Least competition destinations are the target grid cells with the most open canopy, measured as the lowest total biomass of all living trees, and therefore the least light and water competition for AM seeds or seedlings. Post-fire destinations are the target grid cells that experienced fire most recently within the species’ potential range at the time AM was triggered. Lastly, least-fire destinations are target grid cells that experienced fire least recently within the species’ range, which also generally had the lowest fire probability of the four strategies (Figure S7). Altogether, we had eight AM strategies when accounting for each combination of destination type (MB, PF, LF, and MB) and life stage type (DA, GA), plus a ninth control “strategy” of no action (CT). See S1.2 Assisted migration sub-model in the appendix for further details.

#### (C). Climate sub-model

We used the Environment sub-model of ForClim V3.0 for our Climate sub-model (Rasche et al., 2012). This sub-model calculates basic bioclimatic data needed by the Demographic Vegetation sub-model, namely minimum winter temperature, growing degree days, and drought index, which affect species recruitment, growth, and mortality, as well as fire probability. We calculated the local climate conditions in each cell using raw bioclimatic data from WorldClim. For each cell in our model, we calculated the long-term minimum winter temperature as the minimum among the mean monthly temperatures of December, January and February. We used the mean monthly temperature to calculate the growing degree days. We also used the annual actual evapotranspiration and annual potential evapotranspiration at each specific site to calculate the drought index. For detailed description of this sub-model, see previous research using ForClim (Bugmann, 1996; Bugmann & Solomon, 2000) and S1.3 Climate sub-model in the appendix.

#### (D). Fire sub-model

In each time step, the fire sub-model (Figure 1D) simulates tree mortality from fires that occur based on environmental conditions from the Climate sub-model. The fire sub-model has two parts: the fire regime dynamics and fire-induced mortality. To simulate the fire regime dynamics, we used a fire frequency projection model PC2FM to calculate annual fire probability *AFrP* within each grid cell based on its temperature, precipitation, and elevation (Guyette et al., 2017). In a fire year, the model determines fire-induced mortality probability for each tree based on the fire tolerance class of the species, fire severity, and tree diameter at breast height (*DBH,* in *cm*). We considered two types of fire severity following Busing & Solomon (2006): light fires (ground fires) and severe fires (stand-replacing crown fires). We assumed that 5/6 of fires were light, while 1/6 of fires were severe, which is the mean case for montane region calculated based on data from Busing & Solomon (2006). Once a fire (whether severe or light) occurs in a grid cell at one year, we assumed it would spread over the whole cell, thus will affect all trees in this grid cell. Individual trees living in each grid cell then had an additional probability of mortality within the mortality function of the Demographic sub-model (Busing & Solomon, 2006). To determine each tree’s chance of mortality, we divided 23 tree species into four fire tolerance classes, based on the category given by Busing & Solomon (2006) and fire tolerance data from both Busing & Solomon (2006) and USDA Fire Effects Information System (Cooke et al., 2015). *kFiT_s_* is the fire tolerance class of species *s*, with class 1 being the least tolerant and most likely to be killed by fire. All fire tolerance classes other than class 1 exhibit a decrease in mortality probability with increasing tree size (i.e. *DBH*). See Busing & Solomon (2006) and S1.4 Fire sub-model in the appendix for more details.

### III. Model analysis

We validated our model, given the parameters in Tables 1-2 & S2, by simulating forest dynamics for 1500 years under current climate (see S3 Validation in Appendix). We used the best fit model (with the initial seed banks based on current species distributions and including fire-dependent dynamics; see S3.2 Validation Test Results) as the initial state in 2000. Then we ran the model for 100 years of climate change under each of the eight total combinations of AM strategies for different life cycle stages and target locations as well as the control scenario (no-AM). Climate change simulations began with downscaled climate conditions from the year 2000 and ended in 2100, using CanESM5-ssp245 (a scenario with moderately-high greenhouse gas mitigation) and CanESM5-ssp585 (the business-as-usual greenhouse gas emissions scenario) WorldClim data (Fick & Hijmans, 2017). CanESM provides the averaged projection among different climate projection models (Pierce et al., 2018). Within each scenario, we linearly interpolated yearly climate data between each of 2000, 2040, 2060, 2080, and 2100. We simulated 100 replicates of each of the nine total combinations of AM strategies. For each simulation, we calculated the total biomass and gamma diversity weighted by biomass as forestry-oriented outputs and species-specific biomass as conservation-oriented outputs.

To evaluate how the frequency and duration of AM actions affect the outcomes, we analyzed how a range of values for the minimum years between relocation (*I_c_*) and years per relocation (*I_a_*) determined the outcome for AM under the best-performing strategy (LCGA) among the eight AM strategies explored and the CanESM5-SSP585 climate scenario. Within this scenario, we further analyzed AM intensity in terms of the number of seedlings (*N_sd_*) and number of target sites (*N_gd_*) implemented.

## Results

### Forestry-oriented outcomes

On average, both the total biomass and gamma diversity by biomass decreased over the 100 years of simulated climate change, regardless of management approach (Figure 2). The different AM strategies had a negligible effect on total biomass and gamma diversity. In the higher emission SSP585 scenario, we found slightly greater average values for biomass and gamma diversity for AM of seedlings when compared with AM of seeds, and slightly greater values with the least-competition and minimum-distance strategies when compared with the post-fire and least-fire strategies.

**Figure 2:**
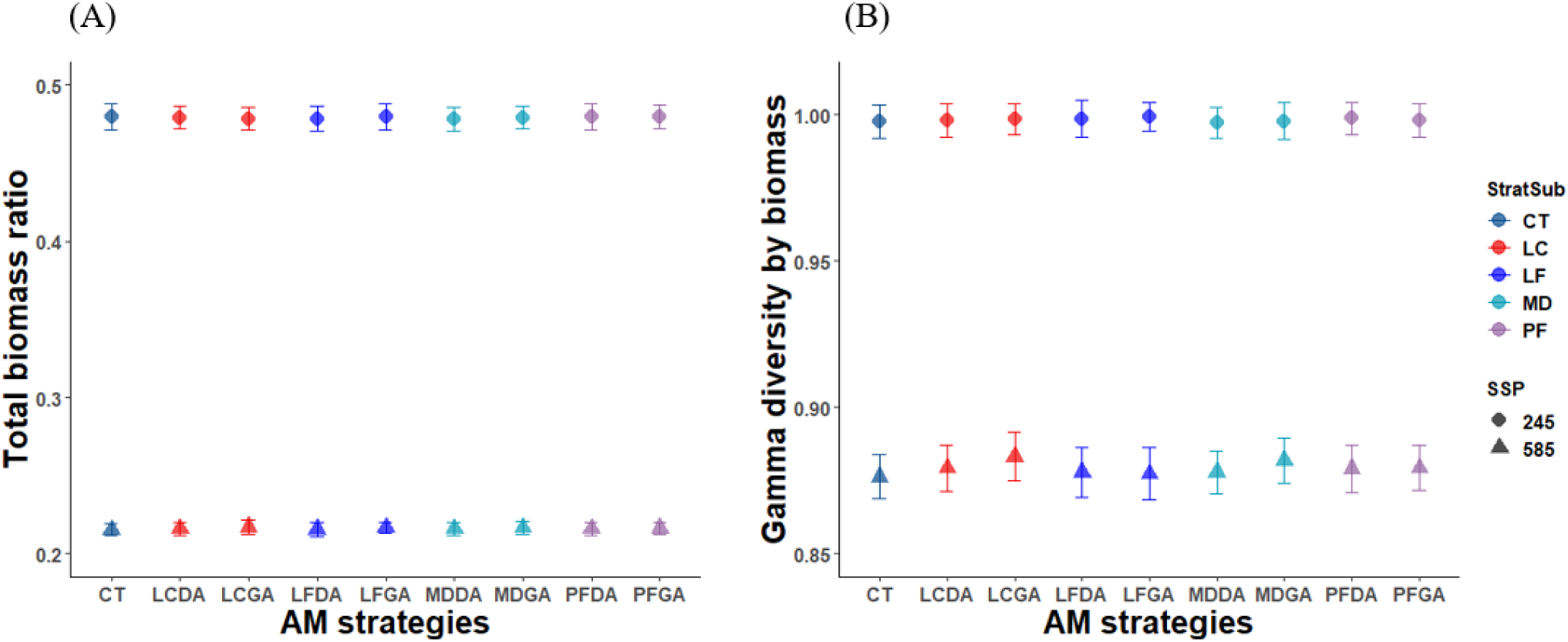
Forestry-oriented outcomes using different AM strategies under two climate change scenarios: SSP245 (moderate climate change; circles) and SSP585 (business-as-usual climate change; triangles). (A) Ratio of total biomass after 100 years of climate change to the total biomass at the initial time step. (B) Gamma diversity by biomass, expressed as the ratio of its value after 100 years to its value at initial time step. The plots show mean values and standard deviation. We classified 8 AM strategies into four groups (LC: least-competition; LF: least-fire; MD: minimum-distance; PF: post-fire) based on their AM destination (different colors), each either introducing seeds (DA) or seedlings (GA), as well as the control group (i.e., no intervention; CT).

### Conservation-oriented outcomes

Under the moderate climate change scenario (SSP245), the biomass of almost all species decreased or had little change compared to the initial state before climate change (Figure S2.A). The single exception was *Acer macrophyllum* (species 5), which has high tolerance to drought, heat, and fire, and whose biomass increased around tenfold (Figure S2.A). Although most species did not frequently fall below the threshold values that trigger AM action, *P. balfouriana* (species 23), one of the least heat-tolerant species as it requires low minimum winter temperature, frequently did, and implementing AM actions increased its biomass (Figure 3A&C).

**Figure 3:**
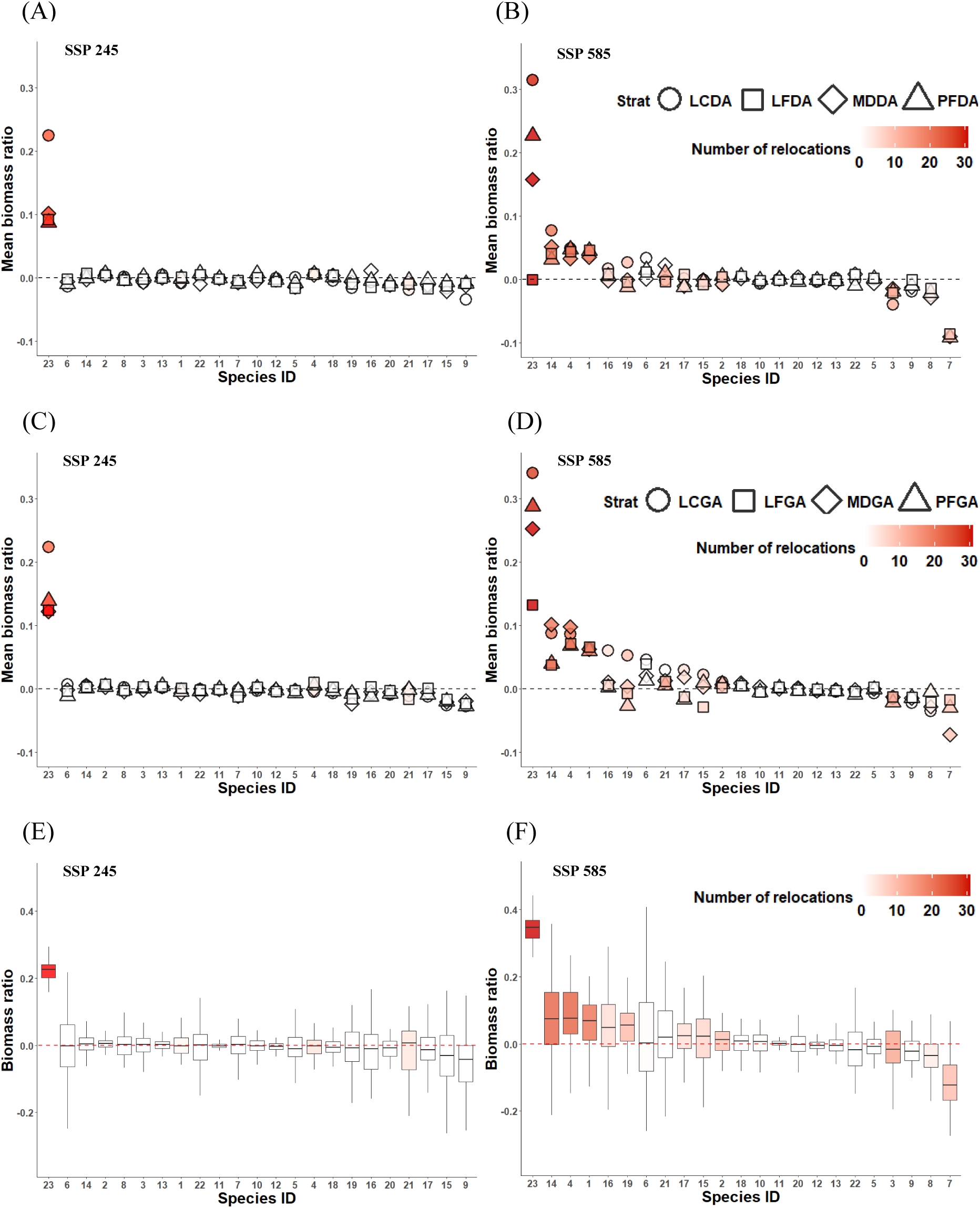
Conservation-oriented outcomes using different AM strategies under two climate change scenarios: SSP245 (moderate climate change; left column) and SSP585 (business-as-usual climate change; right column). In all plots, the biomass ratio is the ratio of the species-specific biomass in 2100 with versus without AM on a log10 scale, and horizontal dashed lines in all panels represent equivalent biomass with versus without AM. In all panels, the color of each dot or bar corresponds number of times AM occurring for that species, while shapes in panels A-D correspond to AM strategies. (A-B) Mean biomass ratio, averaged over 100 repetitions of each AM scenario, for each species given AM at the seed stage. (C-D) Mean biomass ratio for each species given AM at the seedling stage. (E-F) Boxplot of the species-specific biomass ratio under the least-competition seedling AM strategy (LCGA), which is the generally optimal AM strategy to increase species-specific biomass compared with no action.

Under the business-as-usual climate change scenario (SSP585), most species experienced substantial (>70%) biomass decline. Eight of them (*Abies amabilis*, *Abies grandis*, *Abies lasiocarpa*, *Abies procea*, *Chamaecyparis nootkatensis*, *Tsuga heterophylla*, *Pinus albicaulis* and *P. balfouriana*) had a high risk of local extinction, as their biomass decreased by over 90% in our model (Figure S2). They also frequently fell below the threshold values that triggered AM actions (30% of initial biomass; Figure 3.B). Based on a principal component analysis (Figure 4), 7 out of these 8 species (except *P. balfouriana*) were relatively drought-intolerant (low kDrTol), fire-intolerant (low kFi), and cold-adapted species (low kDDMin, kWiTN and kWiTX). The PCA indicated species with strong or intermediate drought and fire tolerance but low tolerance to high temperatures, such as *Pinus jeffreyi* and *P. balfouriana*, were also likely to decline to population sizes that triggered AM (species 19 and 23, Figure 4). By contrast, *A. macrophyllum* (species 5) benefited from future climate change (tenfold increase in biomass), with its high drought tolerance, intermediate fire tolerance, and high heat tolerance (Figure 4 & S2). In short, species with poor adaptation to drought, fire, or warmer conditions constitute the candidate species most likely to go locally extinct without AM (Figure 4, S7 & S9).

**Figure 4:**
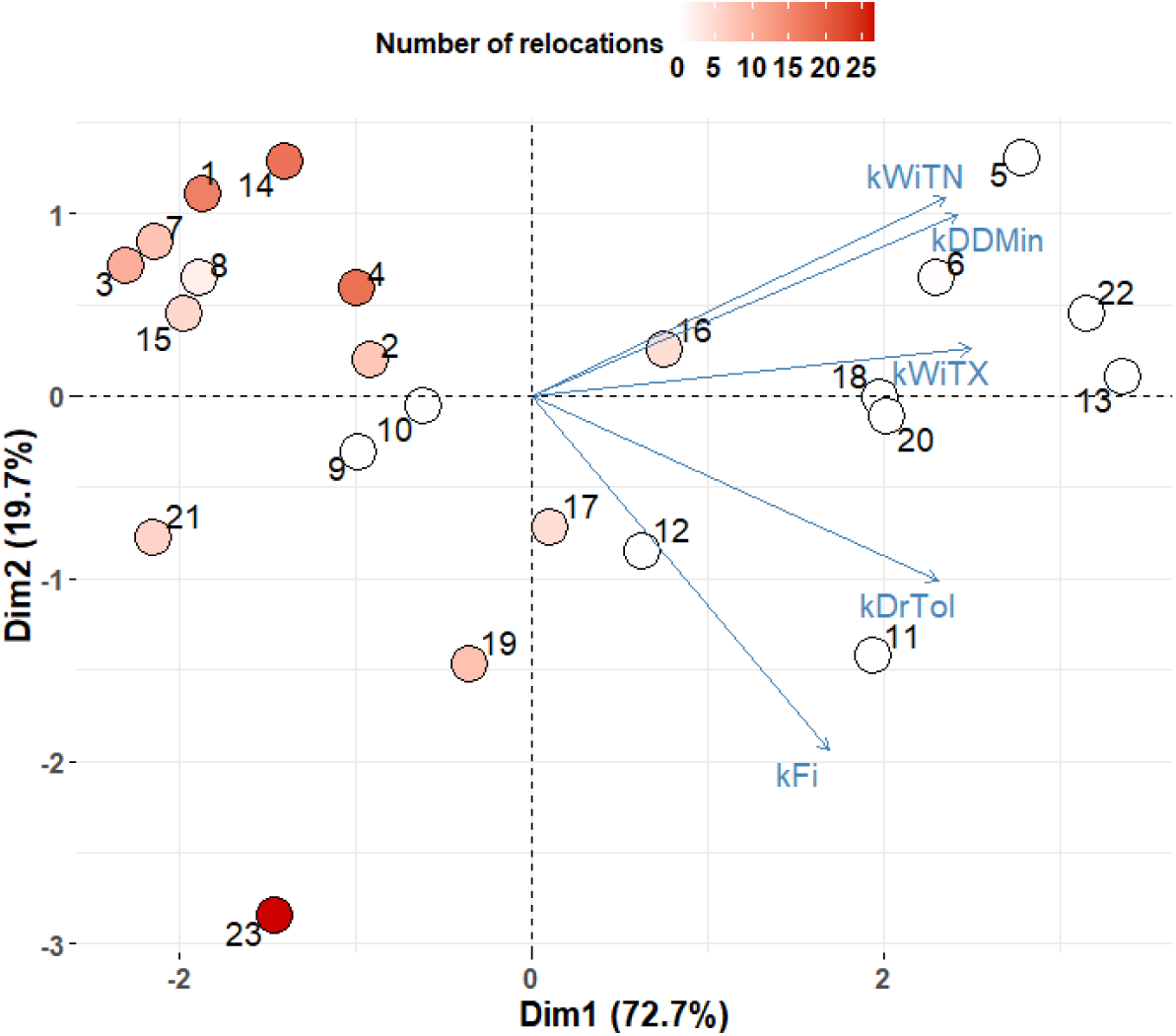
Principal component analysis of species-specific climatic-tolerance parameters. Colors of each dot in this figure correspond to mean species-specific number of relocations under LCGA averaged over 100 repetitions, while numbers correspond to tree species identity. The number of relocations is reflective of the risk of local extinction, namely the more AM actions are needed, the larger the risk. kDDMin: Minimum growing degree-day requirement. kDrTol: Drought tolerance. kFi: Fire tolerance. kWiTN and kWiTX: Minimum and maximum winter temperature tolerances.

Throughout all of our AM simulations, relocations occurred for 15 out of 23 species at least once (Figure 3B). However, only a small number of species consistently benefited substantially from AM (Figure 3B&D). That is, only *P. balfouriana*, *T. heterophylla*, *A. procea* and *A. amabilis* consistently had a higher biomass in simulations that implement AM than those with no action (Figure 3B), with *P. balfouriana* benefiting the most from AM. In contrast, *A. lasiocarpa*, *C. nootkatensis*, *Picea engelmannii* and *Pinus contorta latifolia* had a lower biomass under at least one AM strategy when compared with no action. Among the eight AM strategies, least-competition seedling AM typically led to the greatest average increase in species-specific biomass ratio across all species, though least-competition seed AM and minimum-distance seedling AM performed similarly.

### AM duration and amount

In the analysis of different AM durations, both years per relocation and minimum years between relocation had no prominent effects on the total biomass ratio and gamma diversity by biomass (see Figure S8 in the Appendix). More consecutive years per relocation and fewer years between relocations increased the biomass ratio of *P. balfouriana* (Figure 5C), *Arbutus menziesii*, *Abies concolor*, *Abies magnifica*, and *P. jeffreyi* (Figure S5). AM strategies with more seedlings and more destinations increased biomass ratio of *P. balfouriana* (Figure 5.F), *A. amabilis*, *A. grandis*, *A. procea*, *A. concolor*, *A. magnifica* and *P. jeffreyi* (Figure S6). These species are all AM target species with weak heat, fire or drought tolerance. *C. nootkatensis* had a higher biomass ratio with large number of relocated seedling and fewer destinations, while *P. contorta latifolia* had a higher biomass ratio with fewer seedlings and fewer destinations during AM (Figure S6). These two species, on the other hand, experience biomass decline due to AM-introduced competition (Figure 3).

**Figure 5:**
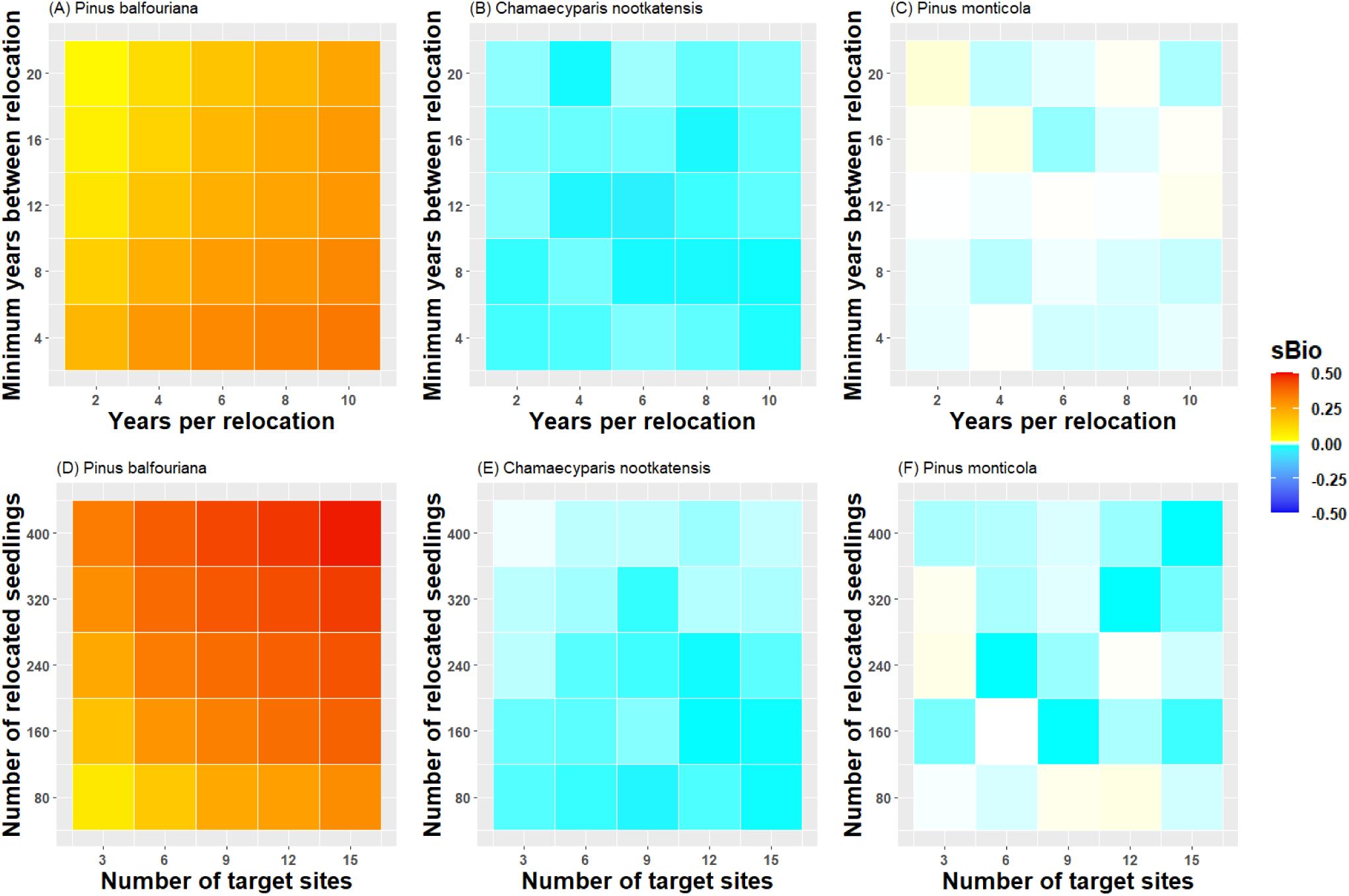
The effect of AM intensity on conservation-oriented outcomes of LCGA under the CanESM5-SSP585 scenario. (A-C) sBio color maps indicate the ratio of species-specific biomass in 2100 using the least competition seeding (LCGA) strategy versus without AM. The impact of years per relocation and minimum years between relocation the sBio of three AM target species: *Pinus balfouriana* (a species with strong tolerance to fire and drought, but weak tolerance to heat), *Chamaecyparis nootkatensis* (a species with weak tolerance to heat, fire and drought) and *Pinus monticola* (a species with moderate tolerance to heat, fire and drought). (D-F) The influence of number of relocated seedling and number of target sites on sBio of the three species.

## Discussion

In general, our simulations suggest that assisted migration can promote conservation goals but would have little effect on forestry goals under climate change. Climate change incurs substantial ecosystem-level and species-level risks in our model in line with previous predictions. On the ecosystem level, our simulations predicted that forest ecosystems in mountains in the western United States are threatened by climate change, where the total biomass of studied tree species declined dramatically under both moderate (reduced by ∼50%) and business-as-usual (by ∼75%) climate change scenarios. Consistent to our results, Loarie et al. (2008) predicted (with a Maxent model) that ranges of about 66% of 591 endemic plant species in California and southern Oregon will decrease by over 80% within this century due to climate change. Similarly, based on simulations of dynamic general vegetation models, Lenihan et al. (2008) and Rogers et al. (2011) projected that California forests might experience notable net losses of carbon by 2100, depending on the climate scenario. In contrast, Zhu et al. (2018) predicted that forest aboveground biomass in Sierra Nevada and Cascades could have a ∼40% increase by 2080 under RCP8.5 scenario, due to forest recovery from agricultural land and growth facilitation by warming. Although these mixed projections indicate some uncertainty in future trajectory of forest biomass shifts (Lenihan et al., 2008; Zhu et al., 2018), most suggest a considerable decline in biomass in the future, which demands urgent conservation action.

On the species level, our simulations predicted that 15 species (out of 23) would experience prominent reduction in biomass under the business-as-usual climate change scenario. Many previous studies supported these projections (Coops & Waring, 2011; Loarie et al., 2008; Stanke et al., 2021). For example, Stanke et al. (2021) found that 3 of these species (*A. lasiocarpa*, *P. engelmannii* and *P. contorta latifolia*) declined in population size over the past two decades due to changes on disturbance regimes and climate. Additionally, Coops & Waring (2011) used a decision-tree based SDM to predict that 11 of the species that declined in our simulations (*A. amabilis*, *A. grandis*, *A. lasiocarpa*, *A. procea*, *C. nootkatensis*, *P. engelmannii*, *P. contorta latifolia*, *P. ponderosa*, *T. heterophylla*, *T. mertensiana* and *P. albicaulis*) will experience 50%-75% range reduction in the Northwestern US by 2080 under a business-as-usual climate change scenario, assuming no dispersal.

In our simulations, AM reduced species-level extinction risks for some vulnerable species by increasing their biomass. For example, *P. balfouriana*, a rare endemic species in California, benefited substantially from AM under both moderate and business-as-usual climate change scenarios, with a total biomass 60% and 120% higher than no action scenarios, respectively. AM also helped to preserve the biomass of *T. heterophylla*, *A. procea*, *A. amabilis*, *A. concolor* and *P. jeffreyi* under the business-as-usual scenario. However, the positive effects of AM on these relocated species were relatively small for the whole ecosystem, resulting in only marginal increases on total biomass and gamma diversity calculated from biomass. In addition, four species declined with AM, likely due to competition from other shifting and relocated species as discussed below.

AM had the greatest species-level conservation benefit (a) to fire, drought, and temperature-intolerant species, (b) when implemented at the seedling stage and with the least competition informing target locations, and (c) when implemented with greater intensity in terms of frequency in time, number of relocated individuals, and number of target locations. We discuss each of these aspects of the AM decision-making process below.

### Candidate species for AM

In our model, the species most at risk from climate change, and therefore strong candidate species for AM, tended to have lower tolerance towards drought, fire, or high temperature. Previous SDMs predicted that these species could experience dramatic future decline in biomass and range under the business-as-usual scenario (Coops & Waring, 2011; Stanke et al., 2021). Our dynamic approach differs from these previous statistical efforts by accounting for competition and dispersal, which have the potential to allow natural range shifts to track climate change. However, our simulations suggest that dispersal would occur too slowly relative to the rate of change under the business-as-usual climate change scenario to track climate change (Figure S2), which reinforces results from these previous studies.

In addition to expected high extinction risk with business-as-usual climate change, species with low fire, drought, and temperature tolerance also experienced the greatest benefit from AM in our simulations. One way to extrapolate from this conclusion to identify potential AM candidates in other systems might be to apply a phylogenetic approach to identify clades that are sensitive to one of these three environmental stresses (Niinemets & Valladares, 2006). Another approach might be to implement a trait-based analysis to estimate climatic tolerance for each tree species. For example, thin bark often indicates low fire resistance (Kidd & Varner, 2019; Stevens et al., 2020), so identifying tree species with this and other tolerance-associated traits as potential candidates for AM could help inform decision-making in less data-rich systems.

### Comparison of different approaches for AM

We found that the optimal AM strategy in our model was least-competition seedling AM, likely because this strategy reduces the risks of establishment failure and competition experienced by transplants. This result confirms the consensus among foresters that matching seedling planting with management of competing vegetation ensures the highest probability of success, in forest habitats all over the world (Brancalion et al., 2019; Nyland, 2001). Survival and growth of tree seedlings are generally negatively associated with surrounding vegetation cover (due to competition for light and other resources) and positively associated with canopy openness (Berkowitz et al., 1995; Duclos et al., 2013; Gerhardt, 1996). Least-competition AM strategies (LCGA and LCDA) targeted grid cells with lower total biomass and therefore higher canopy openness, which have more available light for relocated seeds or seedlings, benefiting the growth of seeds and seedlings after relocation. Additionally, compared to seeds, relocating cultivated seedlings increases the chance of transplants establishing under abiotic filters and light competition.

In addition, both minimum-distance AM and least-competition AM generally led to greater individual-level biomass than post-fire and least-fire AM. The poorer conservation performance of the post-fire strategy is likely because most of the AM target species were drought-and fire-intolerant species (Figure 4), and post-fire AM strategies (PFGA and PFDA) were likely to move individuals to areas with higher fire probability (post-fire target sites have the highest median fire probability among all AM strategies, see Figure S7). The poorer conservation performance of the least-fire strategy is likely because grid cells with the least fire usually developed dense canopies (i.e., generally have higher biomass, see Figure S7), which limits light availability for AM target species under the least-fire AM strategies (LFGA and LFDA). The relatively stronger conservation performance of the minimum-distance AM strategies (MDGA and MDDA) is likely because these strategies were more likely to target grid cells that have similar climatic conditions and community composition to the donor location of target species, which are likely to fit their growth requirement better (Young et al., 2020). Another advantage of the minimum-distance AM strategies, which is not captured in our modeling framework, is their potential to reduce risks of introducing invasive pests or pathogens to the recipient community, because relocation is more likely to occur on scales of natural dispersal, hence the relocated species is more likely to have common evolutionary histories with the targeted community (Wallingford et al., 2020; Williams & Dumroese, 2013).

### Accounting for the risks of AM

Our modeling framework captures the potential for two risks of assisted migration: establishment failure for target species and increased competition with non-target species. Several species, such as *A. grandis*, likely experienced establishment failure during some simulations, as these species had no change in biomass despite AM taking place. This could be because the 2100 climate under business-as-usual climate change scenario (ssp585) in our study region was too harsh for growth and survival of such species, so that moving seedlings only within our study region, they could only be moved to somewhere would not support their establishment. As projected by both our model and some previous studies using SDM, the suitable habitats of most studied tree species will shrink and shift to higher latitudes and elevations in our study region (Coops & Waring, 2011). Within species’ current ranges, environmental conditions might allow those species to survive while also impeding any further establishment of their seedlings (Loarie et al., 2008). Given the costs of AM, identifying which species might not benefit from AM using modeling frameworks such as ours can be as crucial as identifying likely target species.

Secondly, our model assessed the risk of increased competition to non-target or target species, as many species had lower biomass in simulations with AM compared to simulations with no action. For example, in the business-as-usual climate change scenario, *A. lasiocarpa*, *C. nootkatensis*, *P. engelmannii* and *P. contorta latifolia* had lower biomass under all eight AM strategies compared with no action. We ascribe these declines to competition rather than loss to the source population because, in our simulations, all seedling cultivation was external as might occur in a greenhouse or nursery (and these four species underwent moderate-to-no AM intervention in our simulations). All four of these species are mainly found within the northern end of our study region in the Cascade Range. Note that this is the northern limit of our study region due to an international border rather than ecological feasibility (see the “Model assumptions” subsection below). A classic theoretical expectation is that competition determines the lower (or equatorial) limit of a species range and abiotic environmental conditions set the upper (or poleward) limit of a species range (Darwin, 1859; McQuillan & Rice, 2015). Though competition from poleward species might limit the ability for weaker competitors to track climate change (Urban et al., 2012), these four species in the northern part of our study region might face increased competition from equatorward competitors shifting poleward (Gilman et al., 2010), and relocation of competitors with AM could intensify that competition. The combination of establishment failure in AM destination and introduced competition in their current range could explain the decrease in biomass of the four species under AM. Alternate management actions for these species might include cross-boundary coordination, including international coordination, for species whose anticipated range shifts span state or national boundaries (Schwartz et al., 2012; Vitt et al., 2010).

### The effects of AM LCGA intensity on risks and benefits

In simulations with higher intensity AM (i.e., shorter cooldown period, longer action period, more relocated seedlings and/or more targeted grid cells), most of the AM target species had a higher final biomass. However, higher AM intensity also increased risks to species that do not benefit from AM. For example, the two species that had lower biomass in AM simulations compared to no action (*C. nootkatensis* and *P. contorta latifolia*) had even lower biomass with greater AM intensity, due to increased competition, especially with a greater number of AM locations. Increased AM intensity might also increase risks not accounted for in our model, such as relocated pests and pathogens, as well as financial costs. Such costs could include logistical support like employee wages, land purchase, seed storage. and seedling cultivation (Pedlar et al., 2012). Overall, increasing AM could inevitably increase the potential for both risks and benefits, while the balance of risks and benefits depends on local management and stakeholder goals and resources (Schwartz et al., 2012).

### Model assumptions

As with any model, our model entails an array of simplifying assumptions. First, the highest resolution of our climatic data is 1km, whereas microclimatic conditions that affect tree demography can vary at much finer spatial scales, e.g., 10 meters (De Frenne et al., 2019; Fick & Hijmans, 2017). Such conditions might further interact with ecosystem dynamics (De Frenne et al., 2021; Lloret et al., 2012). For example, using data from 98 sites across 5 continents, De Frenne et al. (2019) found that forest canopies can cool the understory when air temperatures are hot and warm the understory when air temperatures are cold, buffering the warming and increasingly variable macroclimate. Given this localized variability and buffering, our model might overestimate the negative effects of climate change on seedling/sapling survival and growth and therefore forest biomass and range loss under future climate change. In addition, ignoring the buffering capacity of canopy might underestimate the risks of least-competition AM, which relocates species to open canopy where stronger climatic variability might occur (Dobrowski et al., 2015).

Secondly, we did not simulate the effects of pests or pathogens, which can play an important role in forest dynamics and climate change effects (Bentz et al., 2010), due to the lack of data. Bark beetles, such as *Dendroctonus brevicomis* and *Dendroctonus ponderosae*, are important forest pests in the NAMCZ, accounting for ∼32% of tree mortality in the region (Berner et al., 2017; Negrón et al., 2009). Their population dynamics are connected to climate conditions and water stress, with higher beetle outbreak risks in warmer and drier years and in denser forest stands, which might substantially increase forest tree mortality in a warmer future (Embrey et al., 2012). The NAMCZ is also characterized by important forest diseases, including white pine blister rust (*Cronartium ribicola*), which infects five-needle pines like *P. lambertiana, P. monticola,* and *P. albicaulis*, and Sudden Oak Death (*Phytophthora ramorum*), which infects a number of oaks (*Quercus* spp.). Without considering the impacts of bark beetles and disease, our model likely underestimates tree mortality and thus the extinction risks of some species. In addition, bark beetle and disease susceptibility might further inform AM target locations, with susceptible species preferentially relocated to locations with low infection risk. Of course, translocation of infected seedlings could also expand the range of the disease in question, another complication we did not incorporate in our modeling.

Thirdly, we set the spatial limits by geographic regions and political jurisdiction rather than ecological properties. Specifically, we restricted our study to the mountains of the westernmost United States, where there were few suitable habitats for some northern range species (e.g., *A. lasiocarpa*) to move within our study region.

Cases where poleward expansion crosses international boundaries might rely on coordinated management across countries (Schwartz et al., 2012; Vitt et al., 2010). Accounting for poleward expansion and coordination with Canada in our case might increase persistence of northern species to support conservation goals. In addition, if we considered species from equatorial regions and coordination with Mexico in our model, we might have expected higher final biomass through dispersal and possible relocation from Mexico into southern California.

Lastly, our model did not consider the potential for phenotypic plasticity or rapid evolution to alter physiological tolerance to the simulated stress, which might reduce extinction risk and AM intervention under climate change. On relatively short (within-generation) time scales, tree species’ phenotypes might change within a certain range via epigenetic, transcriptional, or post-transcriptional regulation to better tolerate changing climate (Chown et al., 2010; Nicotra et al., 2010). For example, drier conditions might trigger some tree species to have increased root-to-shoot ratio, which increases water use efficiency (Nicotra et al., 2010). On longer (across-generation) time scales, tree species might be able to adapt to the changing climate via evolution (Alberto et al., 2013). Part of this evolution depends on locally-adapted genes moving in space, which raises the consideration of assisted gene flow (movement of locally-adapted genes within species ranges) in addition to assisted migration as a management strategy for dispersal-limited species (Aitken & Bemmels, 2016; Young et al., 2020). To explore assisted gene flow using our framework, future studies could include evolutionary dynamics (dynamics of genotypes and corresponding phenotypes for each species) analogous to Kelly & Phillips (2019).

### Future application

Applying our approach in new geographic locations to evaluate conservation performance of AM strategies requires data on species-specific physiological parameters (see Table.S2), which might be available in the literature or the TRY dataset (Kattge et al., 2020), as well as species occurrence data and climatic tolerance trait data. The European Alps are one candidate location for implementing this framework to evaluate AM performance, where these parameters are available for hundreds of tree species (Bugmann, 1994). In addition, certain well-studied (and usually economically important) taxonomic groups like the tree genus *Pinus* would also be good candidates for application of our modeling framework. For more data-poor locations, focusing on our broader take-homes of candidate functional types for AM (species intolerant to fire, drought or heat), and the effects of inter-specific competition on AM risks and benefits can inform management approaches.

## Supporting information

Full supplemental document

## Acknowledgement

We thank Dr. Alvaro G. Gutiérrez and Dr. Harald Bugmann for sharing the physiological parameters of 20 tree species among the 23 studies species with us. This project was funded by the National Science Foundation grant #DEB-1655475 to MLB. All authors have no conflicts of interest to disclose.

## Data availability

The major code and example data to replicate the results in this paper is on the GitHub repository: https://github.com/Yibiaozou/UCDavis_Forest_AM.

