## Supplementary material for "Quantifying the capacity for assisted migration to achieve conservation and forestry goals under climate change": Full supplemental document

### Appendix

**S1 Mathematical details of the full model**

**S1.1 Demographic vegetation sub-model**

In this model, we used diameter at breast height ($DBH$) and tree height ($H$) of each individual tree as two state variables to depict forest dynamics. The following text goes through each dynamic within the tree life cycle, i.e., reproduction, growth and mortality.

**S1.1.1 Tree recruitment with long-distance dispersal and dynamic local seed bank**

In ForClim v.3.0, seed banks of all species are present across all study grid cells (emulating perfect dispersal), and all species have the same fecundity. Two environmental filters regulate tree recruitment from this seed bank (Rasche et al., 2012). First, seedling establishment requires the minimum winter temperature ($uWiT$) in a cell to be greater than a species-specific cold tolerance threshold (${kWiTN}_{s}$) and less than a species-specific chilling requirement (${kWiTX}_{s}$). Second, seedling establishment requires the available light at the forest floor of a patch (depends on the leaf cover of the simulated trees present in the patch) to be greater than a species-specific threshold (*kLy_s_*) (Bugmann, 1996). If both these environmental filters apply, then seedlings of species $s$ establish within a given patch $p$ with a probability $k_{i,j,s}$.

We extend ForClim v.3.0 to allow the local seed bank of each cell to change over time. We represent the local seed bank at a certain time *t* as a three-dimension array $A^{t}$ (130 x 12 x 23, representing the number of north-south latitudinal rows by the number of east-west horizontal columns by the number of species). We initialize $A^{t}$ based on current presence-absence state of all the studied species. If species *s* is present in row $i$ column $j$, then $A_{i,j,s}^{0}=1$, and $A_{i,j,s}^{0}=0$ otherwise.

Species begin each simulation with local seed bank made up of species that are present in post-seed stages, but this seed bank changes as new species reach a cell and reach maturity (whether through natural dispersal or through AM). We simulate dispersal by creating a temporary seed bank following long-distance dispersal by wind using a Wald dispersal kernel $f_{s}(x)$, which represents the probability density of a seed to arrive at a distance *x* (in meters) from the source (G. G. Katul et al., 2005), with $u$ as the mean windspeed and *σ* as the standard deviation of the vertical velocity of the air. Seeds fall from a proportional height ($p)$ of $h_{s}$, the average height of species $s$, at terminal velocity *v_s_* with turbulence coefficient $\kappa$. These values yield the species-specific scale parameter

$$\begin{aligned} \mu_{s}=\frac{ph_{s}u}{\nu_{s}},\#\left( 1 \right) \end{aligned}$$

and shape parameter

$$\begin{aligned} \gamma_{s}=\frac{u\left( ph_{s} \right)^{2}}{2\kappa h_{s}\sigma},\#\left( 2 \right) \end{aligned}$$

for the dispersal kernel. Then the kernel for species *s* is

$$\begin{aligned} f_{s}\left( x \right)=\sqrt{\frac{\gamma_{s}}{2\pi x^{3}}}\exp\left[ -\frac{\gamma\left( x-\mu_{s} \right)^{2}}{2{\mu_{s}}^{2}x} \right].\#\left( 3 \right) \end{aligned}$$

The mean probability for seed of species *s* to disperse from its source *L* x *L* cell to all adjacent cells is

$$\begin{aligned} \alpha_{s}=\frac{1}{L}\times\int_{0}^{L} \int_{L-y}^{2L-y} f_{s}\left( x \right)dxdy,\#\left( 4 \right) \end{aligned}$$

which we numerically integrate in R with the package **pracma** and *L*=10000 m. For all species we modeled, the probability of dispersing further than one cell (over 20 km) was negligible (with an order of magnitude smaller than ${10}^{-27}$). For simplicity, we assumed the probability of dispersing in all four cardinal directions was equal, and we ignored the difference in dispersal probability caused by elevation. Therefore, the mean probability for seed of species *s* to remain in the source cell was $1-\alpha_{s}$ and the probability of dispersing into any certain adjacent cell is $\frac{1}{4}\alpha_{s}$. With these, we constructed the long-distance dispersal matrix for species *s*:

$$\begin{aligned} {DK}_{s}= \left[ \begin{matrix} 0 & \frac{1}{4}\alpha_{s} & 0 \\ \frac{1}{4}\alpha_{s} & 1-\alpha_{s} & \frac{1}{4}\alpha_{s} \\ 0 & \frac{1}{4}\alpha_{s} & 0 \end{matrix} \right].\#\left( 5 \right) \end{aligned}$$

To include the effects of dispersal on seed bank dynamics, we create a temporary seed bank in the form of 3D array $B_{i,j,s}^{t}$ (130 x 12 x 23) with the convolution:

$$\begin{aligned} B_{,,s}^{t}=\text{Conv}\left( A_{,,s}^{t}, {DK}_{s} \right).\#\left( 6 \right) \end{aligned}$$

With the temporary seed bank $B^{t}$, we create the effective local seed bank $C^{t}$ after applying environmental filters:

$$\begin{aligned} C_{i,j,s}^{t}=F_{2,s}\left( F_{1,s}\left( B_{i,j,s}^{t} \right) \right).\#\left( 7 \right) \end{aligned}$$

The filter $F_{1,s}(B_{i,j,s}^{t})$ determines whether species $s$ survives minimum winter temperatures and $F_{2,s}\left( B_{i,j,s}^{t} \right)$ determines whether species $s$ has enough light availability to grow (Bugmann, 1996; Gutiérrez et al., 2016). If species $s$ in the effective seed bank pass through both filters in cell {*i,j*} then $C_{i,j,s}^{t}=B_{i,j,s}^{t}$, otherwise $C_{i,j,s}^{t}=0$. Thus, the probability that species $s$ can establish in cell {*i,j*} is

$$\begin{aligned} k_{i,j,s,t}=\left\{ \begin{aligned} \frac{C_{i,j,s}^{t}}{\sum_{k=1}^{23} C_{i,j,k}^{t}} \text{if}\text{ }\sum_{k=1}^{23} C_{i,j,k}^{t}\neq0 \\ 0 otherwise. \end{aligned} \right.,\#\left( 8 \right) \end{aligned}$$

Once establishment takes place, the number of new-established seedlings for each species that pass all environmental filters, $N_{new}$, ranges from 1 to *N_max_* in each patch (using the default value of *N_max_* as 5 in ForClim v.3.0), following a uniform distribution. The initial *DBH* of all newly established seedlings is 1.27 cm, and we calculated the initial height (*H*, in cm) of each seedling as an approximated function of *DBH* (Rasche et al., 2012)*,* with species-specific maximum height ${kHMax}_{s}$ and parameter denoting initial height growth relative to diameter growth *gS*:

$$\begin{aligned} H=137+\left( {kHMax}_{s}-137 \right)\times\left( 1-e^{\frac{-gS\times DBH}{{kHMax}_{s}-137}} \right).\#\left( 9 \right) \end{aligned}$$

At the end of each year, we update the seed bank with any new species that become present in a grid cell where they were previously absent (i.e. basal area of the target species, ${Bas}_{i,j,s}^{t}$, is larger than the minimum presence threshold, *T_p_*). The species that were already present in the old seed bank $A_{i,j,s}^{t}$ are unchanged, as the seed bank of trees is unlikely to decline under relatively dry condition of Mediterranean-climate Sierra Nevada and Cascade Range (Rajjou & Debeaujon, 2008; Sano et al., 2016). Specifically, we calculated the new local seed bank $A_{i,j,s}^{t+1}$ as:

$\begin{aligned} \boldsymbol{A}_{\boldsymbol{i,j,s}}^{\boldsymbol{t+1}}\boldsymbol{=}\boldsymbol{Bin}\left( \left( \boldsymbol{Bin}\left( \boldsymbol{Bas}_{\boldsymbol{i,j,s}}^{\boldsymbol{t}}\mathbf{>}\boldsymbol{T}_{\boldsymbol{p}} \right)\mathbf{+}\boldsymbol{A}_{\boldsymbol{i,j,s}}^{\boldsymbol{t}} \right)\boldsymbol{\geq}\mathbf{1} \right)\boldsymbol{,\#}\left( \mathbf{10} \right) \end{aligned}$

where *Bin* is a function to transfer logical value “TRUE” to 1 and “FALSE” to 0.

We determined the minimum presence threshold by converting LEMMA maps into presence/absence maps using different values of basal-area presence threshold (from 0.01 to 10 m^2^/ha) and comparing them with presence/absence Little’s maps and finding which threshold match the best (based on Kappa statistic). The best presence threshold is 0.1 m^2^/ha, which gives a Kappa value 0.64.

**S1.1.2 Tree growth**

To calculate volume ($V$) increment of an individual tree of species *s* with species-specific growth rate ${kG}_{s}$, Moore (1989) proposed a carbon budget approach with the equation

$$\begin{aligned} \frac{dV}{dt}=\frac{\Delta\left( {DBH}^{2}\times H \right)}{\Delta t}={kG}_{s}\times{DBH}^{2}\times\left( 1-\frac{H}{{kHMax}_{s}} \right).\#\left( 11 \right) \end{aligned}$$

ForClim replaces volume with $DBH$ to calculate diameter increment with two additional parameters: species specific maximum height above breast height (${kB}_{s}$= ${kHMax}_{s}$-137cm), and an index regulating DBH growth to height growth ($kC$= -*gS*/${kB}_{s}$), such that

$$\begin{aligned} \frac{dDBH}{dt}={kG}_{s}\times DBH\times\frac{1-\left( \frac{H}{{kHMax}_{s}} \right)}{2\times{kHMax}_{s}-{kB}_{s}\times e^{kC\times DBH}\times\left( kC\times DBH+2 \right)}.\#\left( 12 \right) \end{aligned}$$

Based on Linder et al. (1997), *gS* depends on light competition. Specifically, we quantify *gS* as a function of relative light availability at height $H$ of individual tree $i$ ($g{AL}_{H,i}$, in %) with two species-specific parameters, the minimum value of *gS* under low light competition pressure (${kSMin}_{s}$) and an index denoting preference for height growth versus diameter growth under strong light competition (${kE}_{s}$), such that

$$\begin{aligned} gS={kSMin}_{s}+{kE}_{s}\times\left( 1-{gAL}_{H,i} \right).\#\left( 13 \right) \end{aligned}$$

Following Rasche et al. (2012), we calculated ${kSMin}_{s}$ and ${kE}_{s}$ based on their linear relationship to species-specific shade tolerance (${kLa}_{s}$) with the equations

$$\begin{aligned} {kSMin}_{s}=1.3\times{kLa}_{s}+39.5,\#\left( 14 \right) \end{aligned}$$

and

$$\begin{aligned} {kE}_{s}=14\times{kLa}_{s}+13.\#\left( 15 \right) \end{aligned}$$

Linder et al. (1997) assumed that $\Delta H$ is proportional to $\Delta DBH$ such that

$$\begin{aligned} \Delta H=f_{h}\times\Delta DBH.\#\left( 16 \right) \end{aligned}$$

They defined $f_{h}$ as a function that distributes volume growth between $DBH$ and height growth,

$\begin{aligned} f_{h}=gS\times\left( 1-\frac{H-137}{{kHMax}_{s}-137} \right).\#\left( 17 \right) \end{aligned}$

Substituting Eq.(17) into the differential of Eq.(11) gives

$$\begin{aligned} \Delta DBH=\frac{\Delta\left( {DBH}^{2}\times H \right)}{2\times DBH\times H+f_{h}\times{DBH}^{2}}.\#\left( 18 \right) \end{aligned}$$

In this model, three limiting factors of light availability, growing degree-days and soil moisture constrain the tree growth rate from reaching its theoretical optimum.

To quantify the effects of light availability, the model calculated the cumulative leaf area index above the height $H$ of individual tree $i$ ${gCumLA}_{H,i}$ as the sum of foliage weight ($gFolW$) of individual trees with heights above height $H$ of tree $i$ divided by patch size ($kPatchSize$)

$$\begin{aligned} {gCumLA}_{H,i}=\frac{1}{kPatchSize}\times\sum_{j|H_{j}>H_{i}}^{n} \frac{{kC2}_{s}}{{kC1}_{s}}\times{gFolW}_{j},\#\left( 19 \right) \end{aligned}$$

where ${gFolW}_{j}$ is foliage weight of individual tree $j$

$$\begin{aligned} {gFolW}_{j}={kC1}_{s}\times{kA1}_{s}\times{DBH}_{j}^{{kA2}_{s}},\#\left( 20 \right) \end{aligned}$$

and ${kA1}_{s}, {kA2}_{s}, {kC1}_{s}$ and ${kC2}_{s}$ are species-specific allometric parameters.

Then we used Beer’s extinction law (Botkin et al., 1972) to calculate ${gAL}_{H,i}$ as exponentially declining with the cumulative leaf area index above the height $H$ of individual tree $i$ (${gCumLA}_{H,i}$)

$\begin{aligned} {gAL}_{H,i}=e^{-0.25\times{gCumLA}_{H,i}}.\#\left( 21 \right) \end{aligned}$

Based on Bugmann (1994), we quantified ${gL}_{1,i}$ and ${gL}_{9,i}$ (the light response function of the most shade-tolerant tree species, and the most shade-intolerant species) as functions of relative available light at the height of individual tree $i ({gAL}_{H,i})$

$$\begin{aligned} {gL}_{1,i}=1-e^{-4.64\times\left( {gAL}_{H,i}-0.05 \right)},\#\left( 22 \right) \end{aligned}$$

and

$$\begin{aligned} {gL}_{9,i}=2.24\times\left( 1-e^{-1.136\times\left( {gAL}_{H,i}-0.08 \right)} \right).\#\left( 23 \right) \end{aligned}$$

With these, the model calculates the growth limiting factors of available light (${gALGF}_{i}$, individual-specific) as a function of ${gL}_{1,i}$, ${gL}_{9,i}$ and species-specific shade tolerance class (${kLa}_{s}$)

$$\begin{aligned} {gALGF}_{i}=\max\left( {gL}_{1,i}+\left( {kLa}_{s}-1 \right)\times\frac{{gL}_{9,i}-{gL}_{1,i}}{8}, 0 \right),\#\left( 24 \right) \end{aligned}$$

In addition, the model quantifies the limiting effects of growing degree-days (${gDDGF}_{s}$) as a function of annual growing degree-days ($uDD$), species-specific minimum requirement of growing degree-days (${kDDMin}_{s}$), and a parameter ($a$) describing the rate that ${gDDGF}_{s}$ increases with $uDD$

$$\begin{aligned} {gDDGF}_{s}=\max\left( 1-e^{\left( {kDDMin}_{s}-uDD \right)\times a},0 \right),\#\left( 25 \right) \end{aligned}$$

Soil moisture limiting factor (${gSMGF}_{s}$) depends on annual drought stress ($uDrStr$) and species-specific drought tolerance (${kDrtol}_{s}$), such that

$$\begin{aligned} {gSMGF}_{s}=\left( \max\left( 1-\frac{uDrStr}{{kDrtol}_{s}}, 0 \right) \right)^{\frac{1}{2}}.\#\left( 26 \right) \end{aligned}$$

The model computes the total limitation ($gGRF$) as the geometric mean of the three limiting factors (Gutierrez et al., 2016)

$$\begin{aligned} gGRF=\left( {gALGF}_{i}\times{gDDGF}_{s}\times{gSMGF}_{s} \right)^{\frac{1}{3}},\#\left( 27 \right) \end{aligned}$$

Combining Eq.(11), (18) and environmental constraints on growth ($gGRF$) gives the final growth equation in our model

$$\begin{aligned} \frac{dDBH}{dt}={kG}_{s}\times DBH\times\frac{1-\left( \frac{H}{{kHMax}_{s}} \right)}{2\times H+f_{h}\times DBH}\times gGRF.\#\left( 28 \right) \end{aligned}$$

At the end of each time step, we calculated basal area (${gBas}_{i}$, in m^2^/ha) and above ground biomass (${gBio}_{i}$, in kg) for each individual tree $i$ with the equations

$$\begin{aligned} {gBas}_{i}=\frac{{DBH}_{i}^{2}}{4\times kPatchSize},\#\left( 29 \right) \end{aligned}$$

and

$$\begin{aligned} {gBio}_{i}=0.12\times{DBH}_{i}^{2.4}+{gFolW}_{i}.\#\left( 30 \right) \end{aligned}$$

**S1.1.3 Tree mortality**

We considered three types of mortality in our model: baseline, stress-induced, fire-induced. For actively growing trees, it is commonly assumed that no more than ~2% of the trees can reach their maximum age (${kAMax}_{s}$) when there is only baseline mortality (Botkin et al., 1972; Bugmann, 1996). This gives the baseline mortality probability as

$$\begin{aligned} {gPm}_{1}=\frac{4}{{kAMax}_{s}}.\#\left( 31 \right) \end{aligned}$$

The stress-induced mortality rate applies when the increment of a tree’s $DBH$ falls below 10% of the expected optimal $DBH$ increment due to stress, set as

$$\begin{aligned} {gPm}_{2}=\left\{ \begin{aligned} 0.123, \text{stress applies} \\ 0, \text{ otherwise} \end{aligned} \right..\#\left( 32 \right) \end{aligned}$$

When a grid cell is on fire, the fire-induced mortality rate depends on the species-specific fire tolerance class (${kFiT}_{s}$) and declines exponentially with the individual tree’s ${DBH}_{i}$ (Busing & Solomon, 2006) such that

$$\begin{aligned} {gPm}_{3}=\left\{ \begin{aligned} 1, {\text{if} kFiT}_{s}=1 \\ e^{\left( \left( -\left( 1-gFsev \right)\times0.00202-0.00053 \right)\times{DBH}_{i} \right)},{i\text{f} kFiT}_{s}=2 \\ e^{\left( \left( -\left( 1-gFsev \right)\times0.02745-0.00255 \right)\times{DBH}_{i} \right)},{\text{if} kFiT}_{s}=3 \\ e^{\left( -0.00053\times{DBH}_{i} \right)}-0.5-\left( 1-gFsev \right)\times0.5,{\text{if} kFiT}_{s}=4 \end{aligned}. \right. \#\left( 33 \right) \end{aligned}$$

These three mortality rates act on a tree independently, i.e. a tree dies if any one of them applies.

S1.2 Assisted migration sub-model

In this sub-model, we simulated the relocation of seeds or seedlings of a species to a new location if that species’ relative biomass to its initial biomass falls below a threshold value $T_{AM}$. During any timestep when a population of any given species falls below the threshold, we simulated an AM action for this species. Each AM action repeats for $I_{a}$ years (years per relocation), followed by a cooldown period with a minimum of $I_{c}$ years between relocation event such that there is no AM on this species (Backus & Baskett, 2021). We tried multiple values for $T_{AM}$ (0.2, 0.25, 0.3, 0.35, 0.4), and they all gave analogous results. Therefore, we use $T_{AM}$=0.3.

To identify target sites for each target species, we selected grid cells where the projected growing degree-days, drought stress and minimum winter temperature in 2100 were within the species’ tolerance capacity to these bioclimatic conditions (${kDDMin}_{s}$, ${kDrtol}_{s}$, ${kWiTN}_{s}$ and ${kWiTX}_{s}$). During relocation years, the number of target grid cells for each AM action is *N_gc_*.

We simulated AM of two life stages: seed and seedling. We simulated seed assisted migration (DA) by modifying the seed bank (relative recruitment) of the AM target species in the target cell {*i,j*}, assuming the target species *s* has a 30-times recruitment advantage, i.e. $B_{i,j,s}^{t}=30$ in Eq.(7). For seedling assisted migration (GA), we based our simulation on the common practice in forestry of cultivating seedlings in greenhouse for several years (Haase et al., 2019) before moving them to a target location. We simulated this by directly moving a certain number of seedlings (*N_sd_* seedlings per 200 patches in a grid cell) of the target species with *DBH* = 1.27 *cm* (average size of cultivated seedlings among different tree species; (Sáenz-Romero et al., 2021)) into a target grid cell, assuming a constantly-available supply. During each year of AM action, this sub-model simulates seed or seedling AM on all *N_gc_* target grid cells.

Within the potential target range determined by the anticipated future suitable climate, we considered four types of destination site-selection for AM: minimum-distance destinations (MD), least-competition destinations (LC), post-fire destinations (PF) and least-fire destinations (LF).

We depicted the set of grid cells in current range of species $s$ as ${kCurRange}_{s}$, while the set of gird cells in its potential 2100 range as ${kTarRange}_{s}$. The model selected minimum-distance destinations (${kTarCell}_{MD}$) as the target grid cells within the species’ potential 2100 range that are closest to the current range of the target species,

$$\begin{aligned} {kTarCell}_{MD}=\min_{Cell\in{kTarRange}_{s}} \left( \text{Distance}\left( \text{Cell}, {kCurRange}_{s} \right) \right),\#\left( 34 \right) \end{aligned}$$

where $Distance$ is a function that calculates Euclidean distance between two grid cells.

Least competition destinations (${kTarCell}_{LC}$) were the grid cells within potential future range *kTarRange_s_* with the most open canopy so there would be less light competition for AM seeds/seedlings,

$$\begin{aligned} {kTarCell}_{LC}=\min_{Cell\in{kTarRange}_{s}} \left( \text{Biomass}\left( \text{Cell} \right) \right).\#\left( 35 \right) \end{aligned}$$

Post-fire destinations (${kTarCell}_{PF}$) were the grid cells within potential future range *kTarRange_s_* that experienced fire most recently within the species’ potential range at the AM year,

$\begin{aligned} {kTarCell}_{PF}=\min_{Cell\in{kTarRange}_{s}} \left( IntervalLastFire\left( \text{Cell} \right) \right).\#\left( 36 \right) \end{aligned}$Here, $IntervalLastFire$ is a function that gives the number of years between the last fire year and current time step.

Lastly, least-fire destinations (${kTarCell}_{LF}$) were grid cells within potential future range *kTarRange_s_* that experienced fire least recently within the species’ range,

$$\begin{aligned} {kTarCell}_{LF}=\max_{Cell\in{kTarRange}_{s}} \left( IntervalLastFire\left( \text{Cell} \right) \right).\#\left( 37 \right) \end{aligned}$$

Altogether, we had eight AM strategies when accounting for each combination of destination type (MB, PF, LF, and MB) and life stage type (DA, GA), plus a nineth control “strategy” of no action (CT).

**S1.3 Climate sub-model**

This sub-model generates annual growing degree-days ($uDD$), annual drought stress ($uDrStr$) and minimum winter temperature ($uWiT$) for Demographic Vegetation sub-model from raw climatic data.

We calculate $uDD$ as a function of monthly mean temperature ($T_{m}$) and number of days of month $m$ (${kDays}_{m}$)

$$\begin{aligned} uDD=\sum_{m=Jan}^{Dec} \max\left( T_{m}-5, 0 \right)\times{kDays}_{m},\#\left( 38 \right) \end{aligned}$$

$uDrStr$ as a function of annual actual evapotranspiration ($uAET$) and potential evapotranspiration ($uPET$)

$$\begin{aligned} uDrStr=1-\frac{uAET}{uPET},\#\left( 39 \right) \end{aligned}$$

and $uWiT$ as minimum of mean monthly temperature of December ($T_{Dec}$), January ($T_{Jan}$) and February ($T_{Feb}$)

$$\begin{aligned} uWiT=\min\left( T_{Dec}, T_{Jan},T_{Feb} \right).\#\left( 40 \right) \end{aligned}$$

**S1.4 Fire sub-model**

To simulate fire regime dynamics, we used a fire frequency projection model PC2FM to calculate annual fire probability ($AFrP$) within each grid cell (Guyette et al., 2017) by decomposing wildland fire into two components, one term represents effects of physical chemistry on fire frequency ($ARterm$) and another term estimates fuel concentration and quality ($PTrc$) $\begin{aligned} AFrP=\frac{1}{0.232+2.62\times{10}^{-28}\times ARtⅇrm+52\times PTrc}.\#\left( 41 \right) \end{aligned}$

The model quantifies $ARterm$ as a function of Universal Gas Constant (*R*, 0.00831 ${kJ mol}^{-1} K^{-1}$), activation energy (*E_a_*, 132${kJ mol}^{-1}$), annual mean maximum temperature ($Tmax$, in $K$) and molecular collision frequency ($A_{0}$)

$$\begin{aligned} {ARterm=A}_{0}\times e^{\left( \frac{E_{a}}{R\times Tmax} \right)},\#\left( 42 \right) \end{aligned}$$

where $A_{0}$ depends on mean annual precipitation ($P$, in cm) and partial pressure of oxygen (${ppO}_{2}$)

$$\begin{aligned} A_{0}=\frac{P^{2}}{{ppO}_{2}},\#\left( 43 \right) \end{aligned}$$

and the model estimates ${ppO}_{2}$ as a function of elevation ($ELv$, in km)

$$\begin{aligned} {ppO}_{2}= 0.2095\times e^{-0. 12\times ELv}. \#\left( 44 \right) \end{aligned}$$

In addition, the model quantifies $PTrc$ as

$$\begin{aligned} PTrc=\frac{Tmax}{P^{2}}.\#\left( 45 \right) \end{aligned}$$

We set the ratio between frequency of severe fire to light fires as 1:5, based on empirical data from Busing et al. (2006). Therefore, the annual light fire probability is

$\begin{aligned} {AFrP}_{1} = \frac{5\times AFrP}{6},\#\left( 46 \right) \end{aligned}$the annual severe fire probability is

$$\begin{aligned} {AFrP}_{2} = \frac{AFrP}{6}.\#\left( 47 \right) \end{aligned}$$

and the probability for no fire is 1 – $AFrP$. Once a fire (whether severe or light) occurs in a grid cell at one year, we assumed it would spread over the whole cell, thus would affect all trees in that grid cell.

**S2 Parameterization**

To estimate species-specific climate parameters (see Table 1), we extracted tree distribution range data from digital representations of Little’s Map in North America (Bachelet et al., 2010) and raw climate data from WorldClim at 5-arcmin resolution (Gutierrez et al., 2016). Then we calculated corresponding bioclimatic information for each grid cell within the distribution range of each species. We set the minimum growing degree-day requirement (${kDDMin}_{s}$) for a species as 5% of density distributions of growing degree-day extracted out of the species’ observed locations. Similarly, we set drought tolerance (*kDrtol_s_*) as 95% of density distributions of drought index and minimum and maximum winter temperature tolerances (*kWiTN_s_* and *kWiTX_s_*) as 5% and 95% of the density distributions of minimum winter temperature in a species’ observed locations.

Species-specific average tree height (*h_s_*) and seed terminal falling velocity (*v_s_*) are used to calculate long-distance dispersal kernels. For simplification, we assumed a species’ *h_s_* was 50% of its maximum height. *v_s_* came from previous research literature on measuring seed dispersal indexes (Debain et al., 2003; Greene & Johnson, 1995). Seeds fall from a proportional height $p=0.75$ of $h_{s}$, at terminal velocity *v_s_* with turbulence coefficient $\kappa=0.4$. Monthly windspeed data for each grid cell came from the WorldClim database (Fick & Hijmans, 2017). Throughout our simulations, we used the mean maximum monthly windspeed (normally the windspeed during August to November) across all grid cells, such that $\mu=$5.689 *m/s*. We set the standard deviation of windspeed to $\sigma=$ 0.25 *m/s*, which is the minimum value of *σ* in the sensitivity analysis by Nathan et al. (2011).

We used the fire tolerance classes of most species from a study by Busing and Solomon (2006). For *Abies concolor, Abies magnifica, Calocedrus decurrens, Pinus albicaulis, P. balfouriana, P. jeffreyi, Pinus lambertiana,* and *Quercus kellogii,* we obtained fire tolerance information from USDA Fire Effects Information System (Cooke et al., 2015) and assigned them to corresponding fire tolerance classes based on the category given by Busing & Solomon (2006).

The 10 basic physiological parameters for each species in this model are (Rasche et al., 2011; Rasche et al., 2012): maximum height (*kHMax_s_*), maximum age (*kAMax_s_*), growth rate coefficient (*kG_s_*), establishment threshold of relative light intensity (*kLy_s_*), shade tolerance class (*kLa_s_*), maximum relative reductions on maximum tree height induced by climate stress (*kRedMax_s_*) and four photosynthesis-related parameters (*kA1_s_*, *kA2_s_*, *kC1_s_*, *kC2_s_*). We based these species-specific physiological parameters on previous ForClim models (Bugmann & Solomon, 2000; Gutiérrez et al., 2016) for most species (see Table S2)*.* For *A. concolor, A. magnifica, C. decurrens, P. jeffreyi, P. lambertiana* and *Q. kellogii,* we either directly used the parameters from Urban et al. (2000) or estimated them based on phylogenetic closeness between these 6 species and the rest of the species (Gernandt et al., 2005; Urban et al., 2000). For *P. albicaulis* and *P. balfouriana*, we estimated these parameters using an unpublished dataset provided by Dr. Hugh D. Safford and phylogenetic closeness (Gernandt et al., 2005).

**S3 Validation**

**S3.1 Methods**

We validated the model given two initializations by testing its performance on reproducing current species distribution range and basal area distribution. For each initialization, we simulate a variation of the model starting with bare land and run the model for 1500 1-year time steps under current climate conditions (for both fire-presence and fire-absence scenarios). In the first initialization (Initialization I Validation), we assumed the seed bank of 23 species already present in every grid cell of our study region, which is the default setting of most of DVMs. In the second initialization (Initialization II Validation), we took seed bank history into consideration, running the model using the current observed distribution range of each species to determine the geography of the seed bank.

We determined the validity of model projections by comparing the resulting species presence-absence state of the steady-state forest against the observed range of each species (from Little’s map) using the Kappa statistic (Gutierrez et al., 2016), with the assumption that the distribution of tree species is mainly affected by climate tolerance capacities and interspecific interactions (Bugmann, 1996; Bugmann & Solomon, 2000). Kappa values range from −1 to +1, with values > 0.4 indicating a fair degree of agreement and values > 0.6 indicating a good degree of agreement. Values < 0.2 indicate performance no better than random. Following Gutierrez et al. (2016), we complemented Kappa statistic with calculations of AUC (the area under the receiver operating characteristic curve) and sensitivity (the proportion of observed presences that are predicted as such). Then we compared the resulting species-specific basal area density of the steady-state forest against the observed basal area density distribution (from LEMMA map) and calculated the normalized Root Mean Squared Errors (nRMSE) and coefficients of determination (R-square).

**S3.2 Validation Test Results**

The kappa values in Initialization I Validation were greater than 0.2 for 19 out of 23 species and greater than 0.6 for 2 species under at least one scenario (with-fire or no-fire), indicating a relatively accurate prediction of the model (Figure S1.a). In comparison, kappa values in Initialization II Validation were larger than 0.2 for 23 out of 23, larger than 0.4 for 20 and greater than 0.6 for 17 species under at least one scenario (with-fire or without-fire), indicating stronger performance. Including fire in the model had a mixed effect on kappa values, increasing it for some species and decreasing it for others. A larger R-squared value from the regression analysis showed that fire provided generally more accurate prediction on basal area distribution than without fire for validation Initialization I. A smaller nRMSE value for both methods showed that with-fire scenarios were generally more accurate at predicting basal area distribution than without-fire scenarios (Figure S1.c, Table S1). Overall, this suggests that Initialization II with fire was the most accurate model to generate the initial state for climate change scenarios.

**Table S1: Results of the regression analysis for comparing the model-predicted current basal area density with observed basal area density of each species using two validation methods**. Values without parenthesis represent results under the scenario with fire, whereas values within parenthesis represent results under the no-fire scenario.

| ***Species*** | ***InitializationⅠ*** | | ***Initialization Ⅱ*** | |
| --- | --- | --- | --- | --- |
|  | ***R-square*** | ***nRMSE*** | ***R-square*** | ***nRMSE*** |
| *Abies amabilis* | 0.75 (0.63) | 0.74 (2.15) | 0.73 (0.73) | 1.95 (6.06) |
| *Abies grandis* | 0.67 (0.42) | 0.72 (0.96) | 0.87 (0.77) | 1.47 (0.81) |
| *Abies lasiocarpa* | 0.59 (0.53) | 1.51 (1.77) | 0.5 (0.63) | 1.24 (1.47) |
| *Abies procea* | 0.38 (0.21) | 0.98 (1.47) | 0.61 (0.38) | 0.98 (1.77) |
| *Acer macrophyllum* | 0.02 (0.01) | 1.52 (3.92) | 0.27 (0.07) | 1.27 (6.59) |
| *Arbutus menziesii* | 0.23 (0.05) | 1.73 (17.54) | 0.32 (0.11) | 1.69 (7.92) |
| *Chamaecyparis nootkatensis* | 0.26 (0.14) | 2.12 (2.04) | 0.5 (0.53) | 2.03 (2.10) |
| *Picea engelmannii* | 0.33 (0.15) | 1.31 (1.36) | 0.35 (0.23) | 1.18 (1.4) |
| *Pinus contorta latifolia* | 0.00 (0.13) | 1.55 (1.54) | 0.03 (0.03) | 1.49 (1.5) |
| *Pinus monticola* | 0.11 (0.01) | 3.74 (11.22) | 0.0004 (0.02) | 5.37 (22.51) |
| *Pinus ponderosa* | 0.42 (0.42) | 1.62 (1.09) | 0.61 (0.19) | 4.25 (4.18) |
| *Pseudotsuga menziesii menziesii* | 0.5 (0.00) | 0.97 (1.12) | 0.67 (0.19) | 0.48 (0.84) |
| *Quercus garryana* | 0.08 (0.04) | 2.22 (2.47) | 0.29 (0.02) | 1.96 (2.44) |
| *Tsuga heterophylla* | 0.75 (0.70) | 0.65 (2.39) | 0.71 (0.71) | 1.91 (8.52) |
| *Tsuga mertensiana* | 0.08 (0.01) | 1.47 (1.43) | 0.17 (0.09) | 1.30 (1.95) |
| *Abies concolor* | 0.05 (0.25) | 1.46 (2.44) | 0.60 (0.85) | 1.24 (1.84) |
| *Abies magnifica* | 0.06 (0.14) | 1.53 (1.44) | 0.37 (0.50) | 1.11 (4.59) |
| *Calocedrus decurrens* | 0.13 (0.42) | 1.39 (1.67) | 0.37 (0.50) | 1.11 (4.59) |
| *Pinus jeffreyi* | 0.08 (0.11) | 2.72 (2.32) | 0.44 (0.47) | 2.02 (4.89) |
| *Pinus lambertiana* | 0.00 (0.35) | 1.78 (3.54) | 0.73 (0.73) | 1.95 (6.06) |
| *Pinus albicaulis* | 0.09 (0.15) | 12.73 (21.22) | 0.08 (0.13) | 41.6 (49.81) |
| *Quercus kellogii* | 0.34 (0.19) | 1.91 (1.81) | 0.08 (0.19) | 1.90 (1.68) |
| *Pinus balfouriana* | 0.11 (0.13) | 60.51 (37.13) | 0.57 (0.42) | 46.22 (40.58) |

**Table S2: Species-specific physiological parameter set (other than those shown in Table 1).**

| ***Scientific name*** | ***Acronym*** | | ***Species_id*** | ***kHMax_s_* (cm)** | ***kAMax_s_* (years)** | ***kLa_s_*** | ***kG_s_* (cm/year)** |
| --- | --- | --- | --- | --- | --- | --- | --- |
| *Abies amabilis* | | *ABAM* | 1 | 7500 | 600 | 1 | 340 |
| *Abies grandis* | | *ABGR* | 2 | 7600 | 300 | 3 | 357 |
| *Abies lasiocarpa* | | *ABLA* | 3 | 4000 | 300 | 3 | 359 |
| *Abies procea* | | *ABPR* | 4 | 8500 | 600 | 7 | 363 |
| *Acer macrophyllum* | | *ACMA* | 5 | 2800 | 300 | 3 | 280 |
| *Arbutus menziesii* | | *ARME* | 6 | 3400 | 500 | 3 | 154 |
| *Chamaecyparis nootkatensis* | | *CHNO* | 7 | 5300 | 3500 | 3 | 171 |
| *Picea engelmannii* | | *PIEN* | 8 | 5500 | 600 | 3 | 211 |
| *Pinus contorta latifolia* | | *PICO* | 9 | 4600 | 600 | 9 | 226 |
| *Pinus monticola* | | *PIMO* | 10 | 7500 | 600 | 5 | 359 |
| *Pinus ponderosa* | | *PIPO* | 11 | 8000 | 600 | 7 | 324 |
| *Pseudotsuga menziesii menziesii* | | *PSME* | 12 | 5400 | 700 | 7 | 403 |
| *Quercus garryana* | | *QUGA* | 13 | 3700 | 500 | 5 | 161 |
| *Tsuga heterophylla* | | *TSHE* | 14 | 8000 | 700 | 1 | 351 |
| *Tsuga mertensiana* | | *TSME* | 15 | 4600 | 800 | 1 | 203 |
| *Abies concolor* | | *ABCO* | 16 | 7000 | 500 | 3 | 214 |
| *Abies magnifica* | | *ABMA* | 17 | 7000 | 500 | 3 | 145 |
| *Calocedrus decurrens* | | *CADE* | 18 | 5000 | 550 | 6 | 296 |
| *Pinus jeffreyi* | | *PIJE* | 19 | 6000 | 700 | 7 | 241 |
| *Pinus lambertiana* | | *PILA* | 20 | 7000 | 500 | 6 | 338 |
| *Pinus albicaulis* | | *PIAL* | 21 | 4400 | 635 | 7 | 250 |
| *Quercus kelloggii* | | *QUKE* | 22 | 3700 | 500 | 8 | 202 |
| *Pinus balfouriana* | | *PIBA* | 23 | 2300 | 2500 | 8 | 154 |

**Table S2 (Continue)**

| ***Scientific name*** | ***kLy_s_* (%)** | | ***kA1_s_* (kg/cm)** | | ***kA2_s_*** | | ***kC2_s_* (m^2^/kg)** | | ***kC1_s_* (%)** | | ***kRedMax_s_* (%)** | | ***v_s_* (m/s)** | |
| --- | --- | --- | --- | --- | --- | --- | --- | --- | --- | --- | --- | --- | --- | --- |
| *Abies amabilis* | | 0.08 | | 0.23 | | 1.56 | | 6 | | 0.45 | | 47 | | 1.51 |
| *Abies grandis* | | 0.14 | | 0.23 | | 1.56 | | 6 | | 0.45 | | 31 | | 1.74 |
| *Abies lasiocarpa* | | 0.14 | | 0.23 | | 1.56 | | 6 | | 0.45 | | 55 | | 0.63 |
| *Abies procea* | | 0.22 | | 0.1 | | 1.43 | | 6 | | 0.45 | | 57 | | 0.92 |
| *Acer macrophyllum* | | 0.14 | | 0.08 | | 1.43 | | 12 | | 0.35 | | 49 | | 1.08 |
| *Arbutus menziesii* | | 0.22 | | 0.1 | | 1.43 | | 6 | | 0.45 | | 57 | | 0.55 |
| *Chamaecyparis nootkatensis* | | 0.08 | | 0.23 | | 1.56 | | 6 | | 0.45 | | 48 | | 0.64 |
| *Picea engelmannii* | | 0.14 | | 0.23 | | 1.56 | | 6 | | 0.45 | | 45 | | 0.64 |
| *Pinus contorta latifolia* | | 0.47 | | 0.06 | | 1.7 | | 6 | | 0.45 | | 32 | | 0.81 |
| *Pinus monticola* | | 0.14 | | 0.06 | | 1.7 | | 6 | | 0.45 | | 32 | | 0.93 |
| *Pinus ponderosa* | | 0.47 | | 0.23 | | 1.56 | | 6 | | 0.45 | | 37 | | 0.94 |
| *Pseudotsuga menziesii menziesii* | | 0.22 | | 0.06 | | 1.7 | | 6 | | 0.45 | | 31 | | 0.95 |
| *Quercus garryana* | | 0.47 | | 0.1 | | 1.43 | | 12 | | 0.35 | | 49 | | 2.23 |
| *Tsuga heterophylla* | | 0.08 | | 0.23 | | 1.56 | | 6 | | 0.45 | | 40 | | 0.6 |
| *Tsuga mertensiana* | | 0.08 | | 0.06 | | 1.7 | | 6 | | 0.45 | | 42 | | 0.6 |
| *Abies concolor* | | 0.14 | | 0.23 | | 1.56 | | 6 | | 0.45 | | 50 | | 1.17 |
| *Abies magnifica* | | 0.14 | | 0.23 | | 1.56 | | 6 | | 0.45 | | 50 | | 1.13 |
| *Calocedrus decurrens* | | 0.08 | | 0.23 | | 1.56 | | 6 | | 0.45 | | 48 | | 1.19 |
| *Pinus jeffreyi* | | 0.35 | | 0.17 | | 1.4 | | 6 | | 0.45 | | 35 | | 1.12 |
| *Pinus lambertiana* | | 0.27 | | 0.17 | | 1.4 | | 6 | | 0.45 | | 35 | | 1.21 |
| *Pinus albicaulis* | | 0.35 | | 0.17 | | 1.4 | | 6 | | 0.45 | | 35 | | 1.21 |
| *Quercus kelloggii* | | 0.4 | | 0.1 | | 1.43 | | 12 | | 0.35 | | 49 | | 2.23 |
| *Pinus balfouriana* | | 0.4 | | 0.17 | | 1.4 | | 6 | | 0.45 | | 35 | | 1.21 |

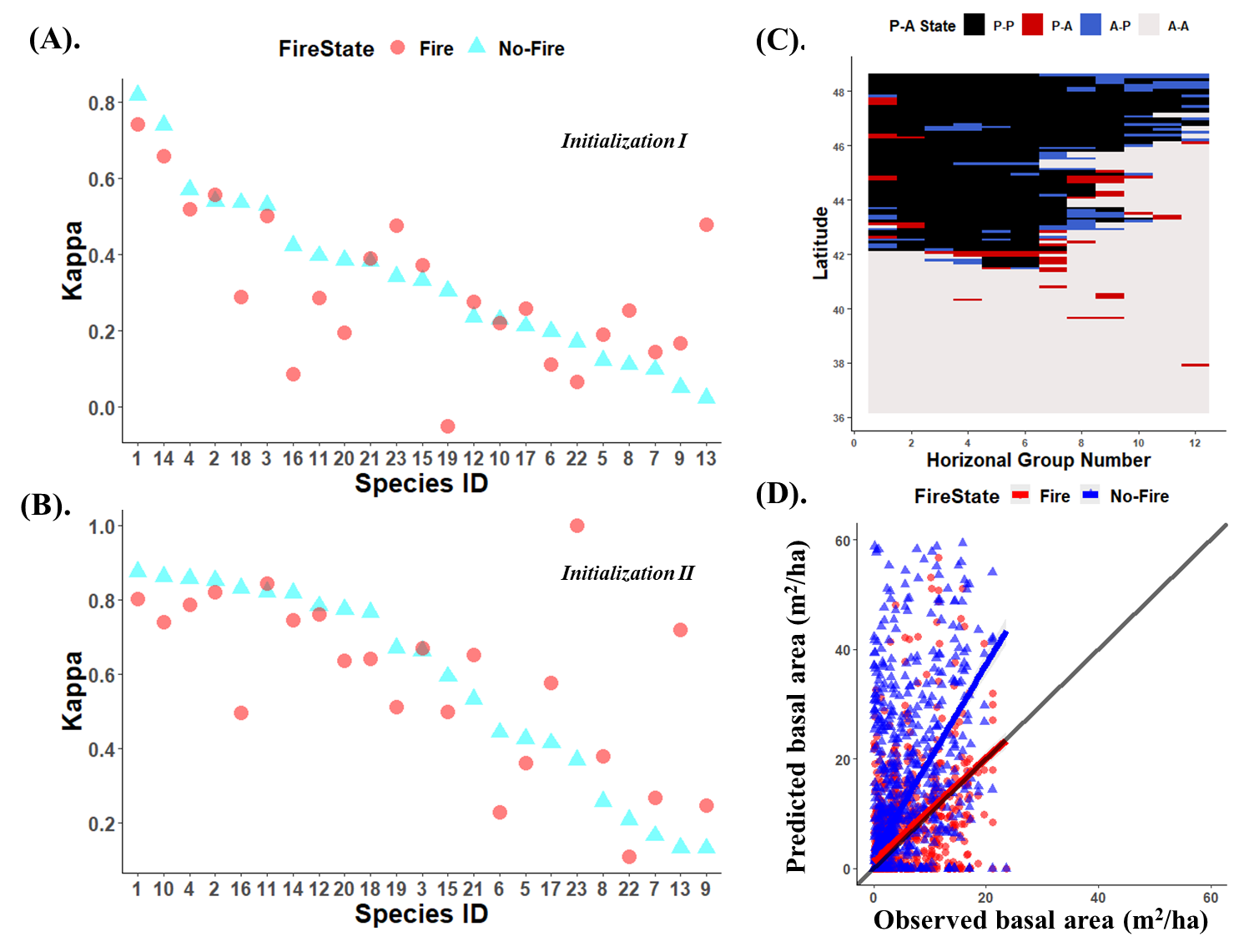

**Figure S1: Results of model validation**. (A-B) Kappa value for model validation given different initial conditions: (A) Initialization Ⅰ with a uniform seed bank and (B) Initialization Ⅱ with a location-dependent seed bank. Red circles represent simulations with fire and blue triangles without fire. Species are ordered in terms of kappa values under the no-fire scenario; see Table 1 for species ID. (C) Map of presence-absence state of species 1 (Abies amabilis, ABAM) under Initialization Ⅰ validation. P-P (black) represents model-predicted presence and observed presence, P-A (red) represents model-predicted presence and observed absence, A-P (blue) is the opposite of P-A, A-A (gray) represents model-predicted & observed absence. (D) Initialization Ⅱ regression results of predicted basal area against observed basal area for species 1, under the with-fire (red) or without-fire scenario (blue). The black solid line represents “x=y”. Higher R-square value indicates better correlation between prediction and observation. The smaller the nRMSE is, the closer those scatters distribute around the solid line, and the better the prediction. For regression results of all species under Initialization Ⅰ & Ⅱ validation simulation, see Appendix Table S1.

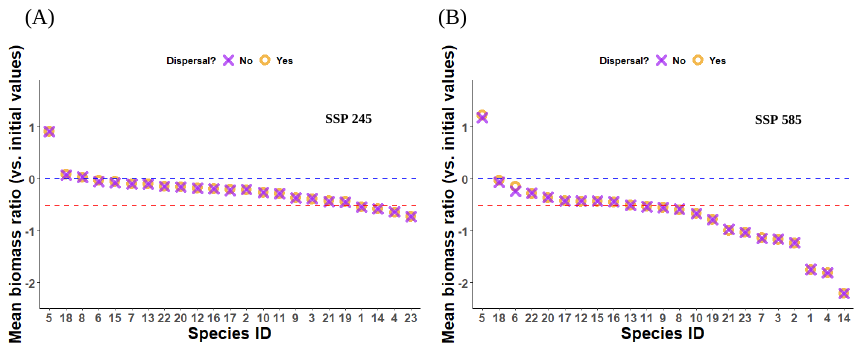

**Figure S2: Conservation-oriented outcomes of biomass for the control group (no AM) in simulations with and without dispersal given SSP245 (A) and SSP585 (B) climate change scenarios**. The mean biomass ratio is the ratio of the species-specific biomass in 2100 with versus without AM on a log10 scale, averaged over 100 repetitions of each AM scenario. Orange circle represents simulations with dispersal, and purple cross represent simulation without dispersal. The species ID on the x axis is ordered by the mean biomass ratio in the scenarios with dispersal.

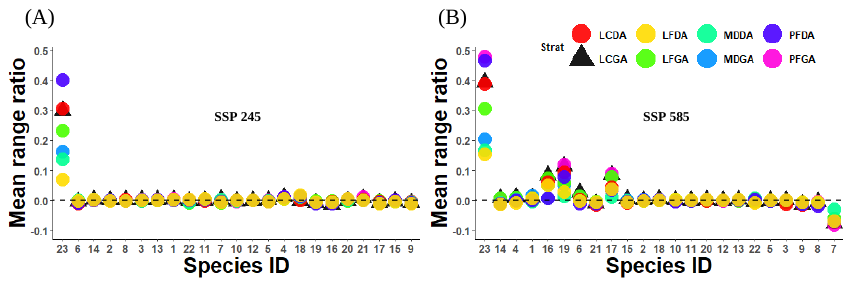

**Figure S3: Conservation-oriented outcomes of range using different AM strategies under SSP245 (A) and SSP585 (B) climate change scenarios**. The mean range ratio is the ratio of the species-specific range in 2100 with versus without AM on a log10 scale, averaged over 100 repetitions of each AM scenario. Colors correspond to different AM strategies.

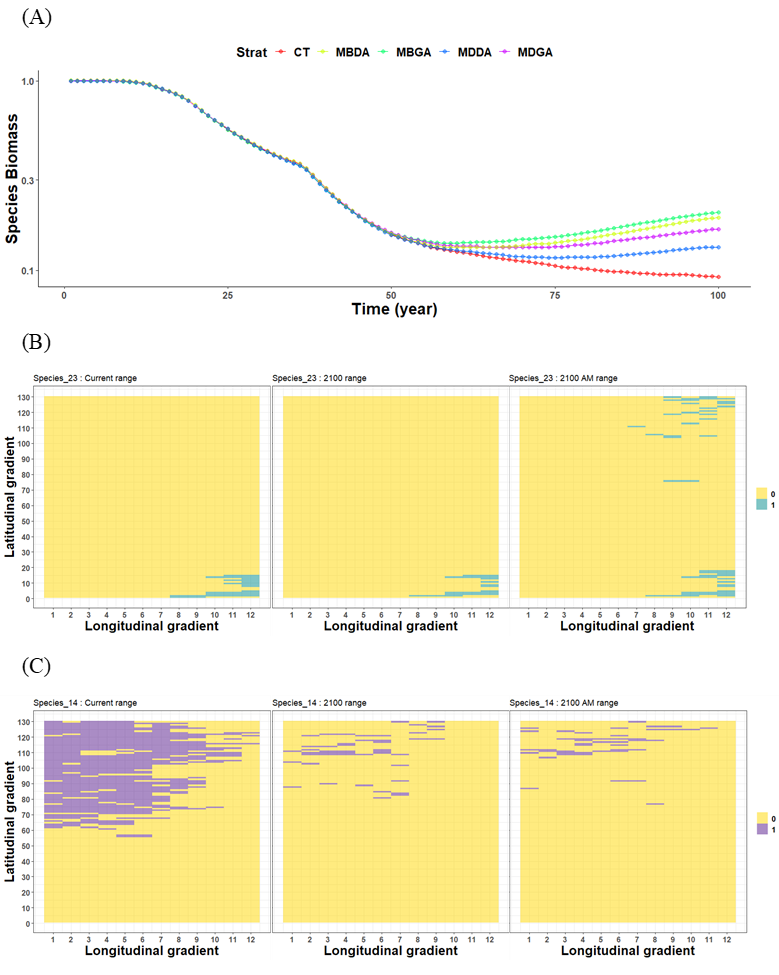

**Figure S4: Spatiotemporal slices for example species.** (A) Relative Biomass (divided by initial biomass) of species 23 (*Pinus balfouriana*) across the 100 years ssp585 climate change under four AM strategies (those that led to the highest biomass) and the control of no intervention (CT). (B) Current range, range in 2100 without AM, and range in 2100 under the least-competition seedling AM (LCGA) strategyof species 23. (C) Current range, range in 2100 without AM, and range in 2100 under LCGA of species 14 (*Tsuga heterophylla*). Subplots (B-C) approximate the whole study region as a spatial matrix with 130 rows along latitude (roughly from 36°11' N to 48°36' N) and 12 columns along longitude (roughly from 122°47' W to 119°3' W).

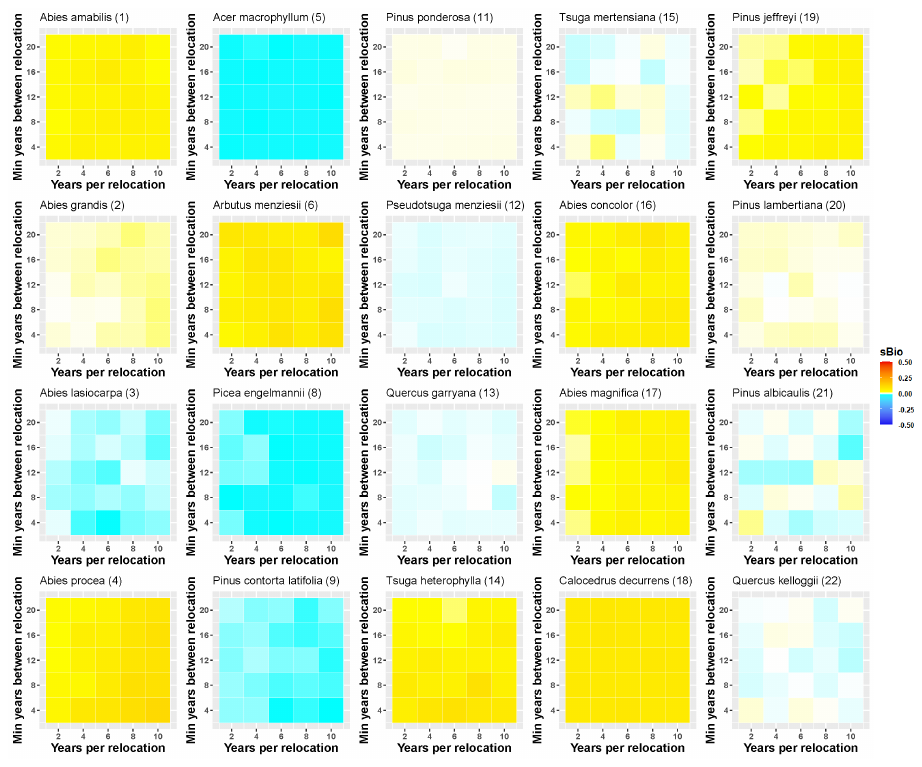

**Figure S5: The impact of years per relocation and minimum years between relocation on the biomass of 20 species under the SSP585 climate change scenario**. The color scale is the same as Fig. 5. See Fig. 5 in the main text for species 7, 10 and 23. Species-specific biomass ratio (sBio) is the biomass of each species in 2100 with LCGA divided by biomass of that species in 2100 without AM.

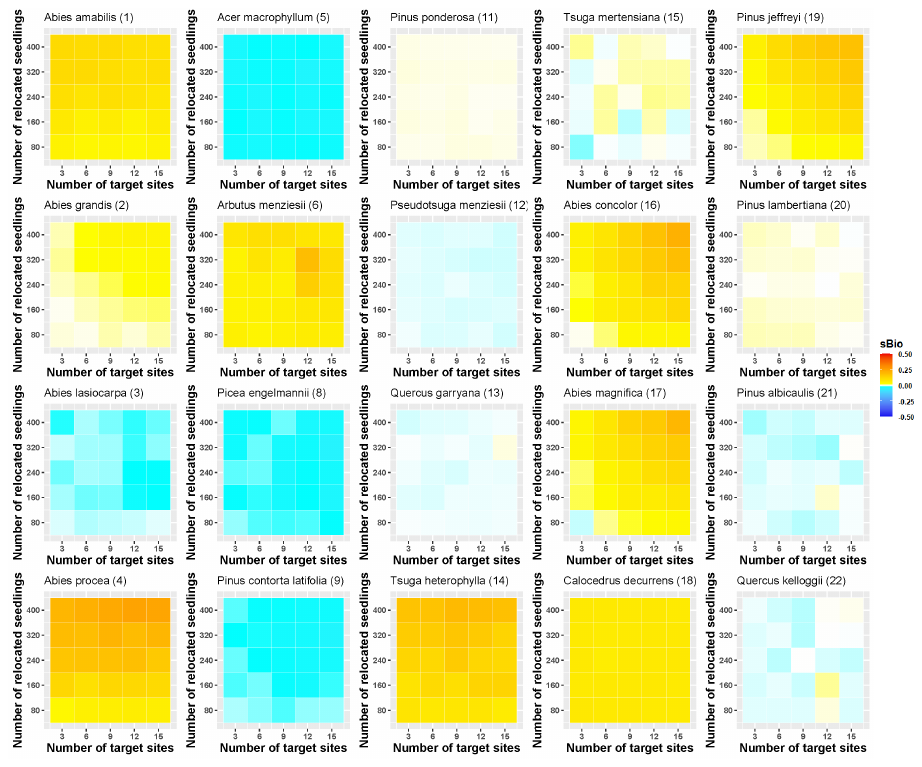

**Figure S6: The impact of the number of relocated seedling and number of target sites on biomass of 20 species under the SSP585 climate scenario.** The color scale is the same as Fig. 5. See Fig. 5 in the main text for species 7, 10 and 23. Species-specific biomass ratio (sBio) is the biomass of each species in 2100 with LCGA divided by biomass of that species in 2100 without AM.

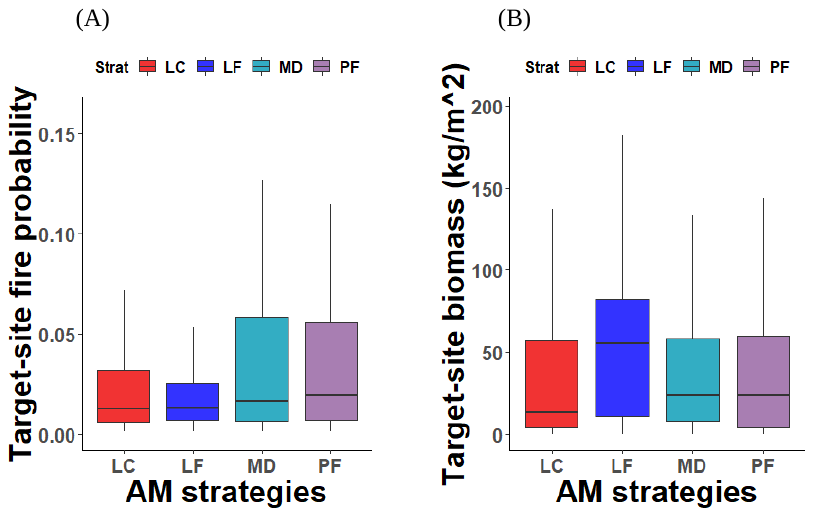

**Figure S7: Target-site properties for different AM strategies.** (A) Distribution of fire probability in AM target sites as it depends on AM strategy. (B) Distribution of the total community biomass in AM target sites as it depends on AM strategy. Color represents different AM strategies. LC: least-competition, LF: least-fire, MD: minimum-distance, PF: post-fire.

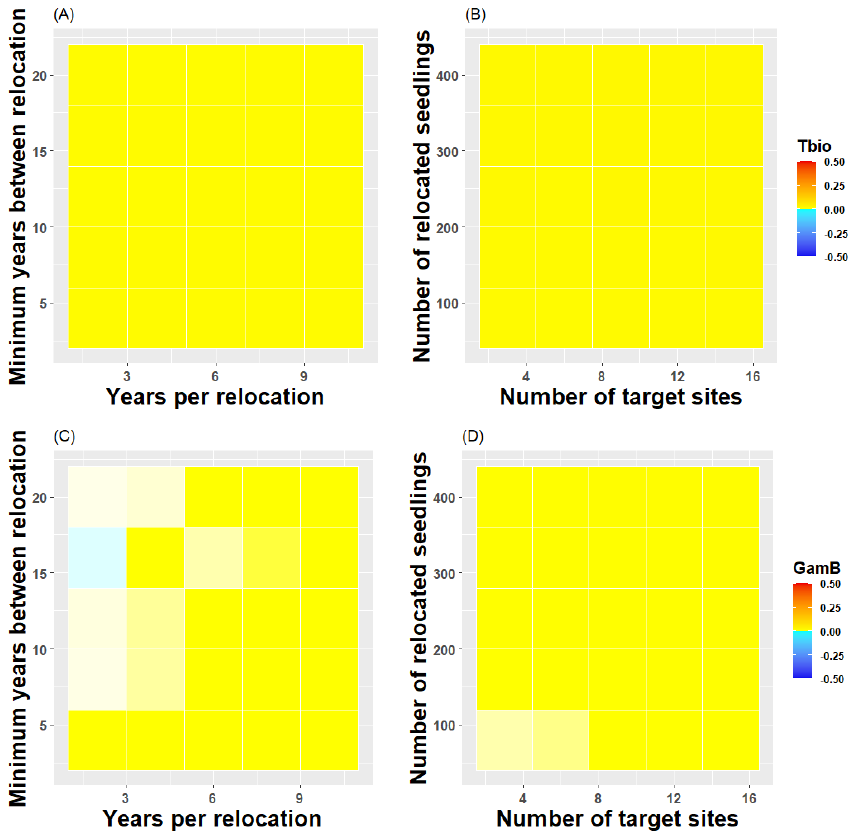

**Figure S8:** **The impact of AM intensity on forestry-goal oriented outcomes:** the impact of minimum years between relocation and years per relocation on (A) total biomass and (C) gamma diversity by biomass, and the impact of the number of relocated seedlings on (B) the number of target sites on total biomass and (D) gamma diversity by biomass, under the SSP585 climate scenario. The color scale is the same as Fig. 5. Both Tbio (total biomass) and Gamma diversity by biomass (GamB) are the ration of their values after 100 years of climate change relative to the initial values.

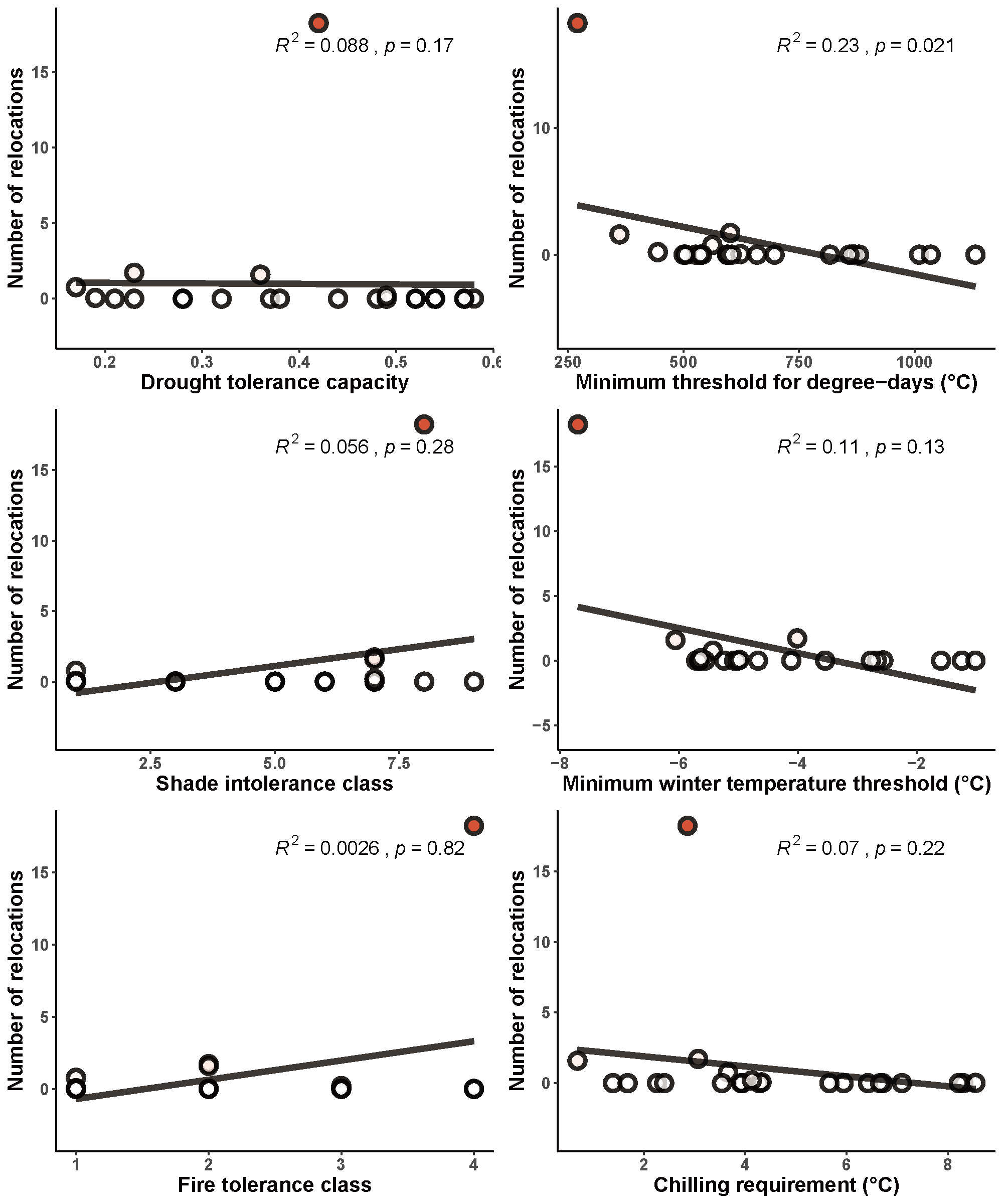

**Figure S9: Pairwise scatter plots of mean number of relocation events versus species-specific climatic tolerance traits.** The color of each point represents the mean number of relocation events with darker colors corresponding to a higher number of relocations. The black solid line indicates the linear regression for each pairwise comparison, with corresponding R2 and p-values at the top of each sub-plot.
